# Mesenchymal stem cell culture within perfusion bioreactors incorporating 3D-printed scaffolds enables improved extracellular vesicle yield with preserved bioactivity

**DOI:** 10.1101/2022.08.30.505860

**Authors:** Stephanie M. Kronstadt, Divya B. Patel, Louis J. Born, Daniel Levy, Max J. Lerman, Bhushan Mahadik, Shannon T. McLoughlin, Arafat Fasuyi, Lauren Fowlkes, Lauren Hoorens Van Heyningen, Amaya Aranda, Sanaz Nourmohammadi Abadchi, Kai-Hua Chang, Angela Ting Wei Hsu, Sameer Bengali, John W. Harmon, John P. Fisher, Steven M. Jay

## Abstract

Extracellular vesicles (EVs) are implicated as promising therapeutics and drug delivery vehicles in various diseases. However, successful clinical translation will depend on development of scalable biomanufacturing approaches, especially due to the documented low levels of intrinsic EV-associated cargo that may necessitate repeated doses to achieve clinical benefit in certain applications. Thus, here we assessed effects of a 3D-printed scaffold-perfusion bioreactor system on the production and bioactivity of EVs secreted from bone marrow-derived mesenchymal stem cells (MSCs), a cell type heavily implicated in generating EVs with therapeutic potential. Our results indicate that perfusion bioreactor culture results in an ~40-80-fold increase, depending on measurement method, in MSC EV production compared to conventional cell culture. Additionally, we demonstrated that MSC EVs generated using the bioreactor system significantly improved wound healing in a diabetic mouse model, with increased CD31+ staining in wound bed tissue compared to animals treated with flask cell culture-generated MSC EVs. Overall, this study establishes a promising solution to major EV translational issues (i.e., scalability and low potency) with potential for adaptation to various EV-based therapeutics and capacity for improvement alongside the continuous advancements in 3D-printing technologies.

## Introduction

Extracellular vesicles (EVs), specifically those from mesenchymal stem cells (MSCs), have emerged as an intriguing therapeutic alternative to whole-cell therapies in a multitude of applications (e.g., sepsis, cancer, wound healing) with additional promise as efficient drug delivery vehicles [1, 2]. With an improved safety profile [3], favorable storage requirements [4], and the ability to traverse biological barriers [5, 6], MSC EVs represent a promising and effective alternative to their parental cells. However, there exist fundamental obstacles that hinder the translation of MSC EVs, including a lack of a rationally designed production platform and low therapeutic potency (i.e., low levels of endogenous EV cargos).

Numerous methods have been employed to attempt to combat these issues, including MSC EV cargo loading and biochemical priming of parental cells [7–9]. These approaches have their merits, but they are generally limited with respect to scalability and could substantially increase production costs and the regulatory burden associated with MSC EV translation. Alternatively, biophysical cues, such as flow-derived shear stress, intrinsically exist in established biomanufacturing set-ups and can be utilized in a highly reproducible and cost-effective manner [10]. Importantly, dynamic culture conditions have been shown to augment stem cell EV production and bioactivity. For example, using GMP-compliant serum- and xeno-free cell culture media, Gobin and colleagues were able to utilize a hollow-fiber bioreactor system to significantly increase MSC EV production across multiple donors while maintaining the functionality of the parental cells [11]. In a separate study, EVs from human dental pulp stem cells (DPSCs) grown in dynamic conditions on a fiber-based scaffold significantly increased axonal sprouting in neurons when compared with DPSC EVs from static conditions [12]. However, in addition to flow-derived shear stress, these systems introduce other cues as the topographical, chemical, and mechanical attributes of the membrane itself can affect cellular behavior, which ultimately convolutes the true effectors at play [13]. Recent studies have utilized simpler systems (e.g., flat-plate bioreactors) [14], allowing a more precise approach to understanding and harnessing flow-derived shear stress but still lack the tunability necessary for dynamic manufacturing and exploratory research needs.

A promising solution exists in 3D-printing technology, which, with the ability to precisely control scaffold geometry and architecture [15], allows for more adaptable systems as well as more precise computational fluid modeling. As the therapeutic profiles of EVs are often shaped by the stress-adaptive responses of their parental cells [16, 17], a tunable platform is vital in the ability to understand these responses and subsequently adjust EV formulations for specific applications. In a previous study, we employed a 3D-printed, biocompatible scaffold coupled with a peristaltic pump, which we termed the bioreactor, to enhance the production of human dermal microvascular endothelial cell (HDMEC) EVs [18]. We also showed that ethanol conditioning-mediated pro-angiogenic effects of HDMEC EVs, previously demonstrated using conventional culture techniques [19], were maintained when cells were cultured in the bioreactor. In the current investigation, we utilized a similar bioreactor to assess the effects of flow-derived shear stress on the production and bioactivity of EVs secreted by bone marrow-derived MSCs as they are a common therapeutic cell source [20]. We specifically focused on wound healing abilities due to the well-documented pro-angiogenic effects of EVs from traditionally-cultured MSCs [21, 22]. Our results suggest that this particular bioreactor set-up can be used to increase MSC EV production while maintaining pro-angiogenic bioactivity, effectively increasing the potency of the EV formulation. Given the tunability of this system, the methods used here have the potential to be applied to various EV-based formulations with the ability to increase effectiveness without sacrificing scalability.

## Methods

### Cell Culture

Bone marrow-derived mesenchymal stem cells (MSCs; passage 2) were purchased from ATCC (PCS-500-012). A kit of MSCs from three different donors was also purchased from RoosterBio to assess donor variability (KT-014). All MSCs were cultured in Dulbecco’s Modification of Eagle’s Medium (DMEM) (Corning, 10-013-CV) supplemented with 10% fetal bovine serum (FBS; VWR, 89510-186), 1% non-essential amino acids (NEAA; Fisher Scientific, 11-140-050), and 1% penicillin-streptomycin (P/S; Corning, 30-002-CI). MSCs were expanded so they were at passage 4 upon seeding into the experiments.

Human embryonic kidney cells (HEK293) were used to assess the effects of bioreactor culture on EV production in a relatively mechanically-inert cell type. HEK293 cells were purchased from ATCC (CRL-1573) and cultured in DMEM supplemented with 10% FBS and 1% P/S. During EV collection, MSC or HEK293 culture media was switched to media containing 10% EV-depleted FBS (referred to as EV-depleted media). EV-depleted FBS was produced by centrifuging heat-inactivated FBS at 118,000 × *g* for 16 h and filtering the supernatant through a 0.2 μm bottle-top filter for later use in media supplementation.

Human umbilical vein endothelial cells (HUVECs) pooled from multiple donors (C-12203) and human dermal microvascular endothelial cells (HDMECs) (C-12212) were obtained from PromoCell and used for the testing of EV angiogenic bioactivity. HUVECs or HDMECs were cultured on tissue culture polystyrene flasks coated with 0.1% gelatin at 37°C for 1 h prior to seeding. HUVECs and HDMECs were cultured in complete endothelial growth medium-2 (EGM2; PromoCell C-22111) supplemented with 1% P/S and used at passages 3-5 in experiments. During experiments, HUVECs and HDMECs were maintained in endothelial basal medium-2 (EBM2; PromoCell, C-22221) supplemented with 0.1 %FBS and 1% P/S.

### 3D-Printed Scaffold Design and Fabrication

The scaffold design involved a series of small pillars that were 1 mm in diameter and situated 2.5 mm apart in a 50 cm^2^ cell growth area within a 12 cm^3^ volume construct. This design was chosen to provide a sufficient culture surface area in a highly-ordered pillared array that was amenable to cell removal and immunofluorescence imaging. Additionally, the pillared array was situated in the middle of the construct to allow for fully-developed flow prior to entering the pillars. The architecture also allowed for media to perfuse the circuit around and in-between the pillars, facilitating active transport of nutrients and gases from gas permeable tubing, and provided a mechanism to control fluid characteristics predictably throughout. Computational fluid modeling was accomplished using the SolidWorks (Dassault Systèmes, Velizy-Villacoublay, France) Flow Simulation add-in. Fluid flow was analyzed at flow rates of 1, 5, and 10 mL/min by modulating the inlet volume rate with surface shear stress, flow profiles, and fluid velocity (computed and recorded).

Scaffold designs were exported into stereolithography (.stl) files following computational design and scaffold models were oriented, fixed, and supported using Magics 18 (Materialise, Leuven, Belgium). Solid objects were fabricated out of a clear, biocompatible, acrylate-based material (E-Shell 300; EnvisionTEC, Inc.) using a commercially available stereolithography apparatus (EnvisionTEC Perfactory 4 Mini Multilens; Gladbeck Germany). Excess resin was removed by submerging printed objects in 99% isopropanol (Pharmco-Aaper, Shelbyville, KY) for 5 min, followed by flowing 99% isopropanol through the scaffolds, and blowing the interior dry with filtered air. The process was repeated until all excess material was removed from the interior of the object. Complete resin curing was achieved with 2000 flashes of broad-spectrum light (Otoflash, EnvisionTEC, Inc.). Scaffolds were cleaned in 100% ethanol (Pharmco-Aaper, Shelbyville, KY) for >30 min to leach any remaining soluble contaminants before proceeding to the sterilization and rehydration steps described below.

### Scaffold Sterilization, Coating, and Cell Seeding

3D-printed scaffolds were submerged in fresh 100% ethanol and sterilized in an ultraviolet sterilizer (Taylor Scientific, 17-1703) for 10 min. Scaffolds were gradually rehydrated by submerging in sterile solutions with a progressively increasing volume of pH 7.4 sterile 1X PBS (i.e., 25:75 PBS:EtOH for 5 min, 50:50 PBS:EtOH for 5 min, 75:50 PBS:EtOH for 5 min) and placed in 100% sterile 1X PBS in the fridge until use. Once ready to use, scaffolds were coated with 0.5 μg/cm^2^ fibronectin in sterile water for 30 min at 37°C. Open ends of the scaffolds were closed using platinum-cured tubing and binder clips to prevent leakage. Scaffolds were drained and passage 4 MSCs were seeded into the scaffolds at a seeding density of 2,500 cell/cm^2^ in 15 mL of EV-depleted MSC media. Cells were allowed to attach for 24 h at 37°C before connecting the scaffold to the bioreactor or replacing the media and keeping the scaffold detached as a static control. In parallel, MSCs were also seeded in 75 cm^2^ tissue culture flasks at the same density as within the scaffolds in EV-depleted MSC media as an additional static control.

### Perfusion Bioreactor Assembly

After initial cell seeding, the 3D-printed scaffolds were connected to a Masterflex L/S Digital Drive (Cole-Parmer) coupled with a pump head to circulate medium at 1, 5, or 10 mL/min. Open ends of the tubing on either end of the scaffold were affixed to a media reservoir with 50 mL of EV-depleted MSC media. The pump head allowed for up to 4 lines to be connected at a time at a single flow rate. The assembly was then placed within a cell culture incubator at 37°C with 5% CO_2_, with the scaffold chamber secured on the incubator sidewall with flow anti-parallel to gravity. The complementary flask and scaffold controls were also incubated at 37°C with 5% CO_2_. Scaffolds were situated vertically to ensure complete submergence in medium.

### Cell Staining and Imaging

After 24 h culture in the bioreactor, MSCs were fixed in the scaffolds with 4% paraformaldehyde (PFA) and 1% sucrose for 15 min and washed three times with 1X PBS. Prior to staining, cells were permeabilized for 5 min with a 300 μM sucrose, 100 μM sodium chloride, 6 μM magnesium chloride, 20 μM HEPES, and 0.5% Triton-X-100 solution. Cellular actin was stained in a 1:100 dilution of AlexaFluor 488 Phalloidin (Invitrogen, A12379) in 1X PBS for 20 min and visualized on a Nikon Ti2 Microscope (Nikon, Minato City, Tokyo, Japan).

### EV Isolation and Characterization

Conditioned media from MSCs cultured in EV-depleted media were collected and underwent a series of differential centrifugation steps which entailed a final centrifugation step of 118,000 × *g* for 2 h as previously described [23, 24]. After resuspending the pelleted EVs in 1X PBS, they were transferred to a Nanosep 300 kDa MWCO spin column (Pall, OD300C35) and centrifuged at 8,000 × *g* until all PBS was removed (~8-12 min). The EVs were washed two more times in a similar fashion. EVs collected at the top of the column were then resuspended in a desired volume of 1X PBS and sterile filtered using 0.2 μm syringe filters. The total surface protein concentration of the washed EVs was measured by BCA. EV size distribution and concentration were evaluated using a NanoSight LM10 (Malvern Instruments; Malvern, UK) with Nanoparticle Tracking Analysis (NTA) software version 2.3. Each sample was measured three times with a camera level set at 14 and acquisition time of 30 s. Approximately, 20-100 objects per frame with more than 200 completed tracks were analyzed for each video. The detection threshold was set at the beginning of each sample and kept constant for each repeat. EVs were also quantified based on the amount of total immunoreactive CD63 within isolated EV samples using the ExoELISA-ULTRA Complete Kit (System Biosciences, EXEL-ULTRA-CD63-1) per the manufacturer’s protocol. Briefly, 10-25μg of EVs (based on BCA analysis) were used for the analysis. The number of EVs was obtained using an exosomal CD63 standard curve calibrated against NTA data. Total number of EVs was then calculated based on resuspension volume and final data was expressed as total # of EVs/cell.

Immunoblotting was then performed to confirm the presence of EVs and purity of each sample. Based on the BCA analysis, 10 μg of protein from each EV sample was used for analysis and compared with cell lysate. The presence of various EV-associated proteins was assessed using primary antibodies for Alix (Abcam, ab186429), TSG101 (Abcam, ab125011), CD9 (Abcam, ab92726), CD63 (Proteintech, 25682-1-AP), actin (Cell Signaling Technology, 4970), and GAPDH (Cell Signaling Technology, 2118), while the absence of contaminating proteins were confirmed using antibodies for calnexin (Cell Signaling Technology, 2679), HSP90 (Abcam, ab13492), and AGO2 (Abcam, ab186733). All primary antibodies were added at a 1:1,000 dilution, except GAPDH which was diluted 1:2,000. A goat anti-rabbit secondary (LI-COR Biosciences, 926-32211) was added at a 1:10,000 dilution. Protein bands were detected using a LI-COR Odyssey CLX Imager. All EVs were used within three days or frozen at −20°C and used within 2 weeks with no more than 1 freeze/thaw cycle.

### Transmission Electron Microscopy

A portion of each EV sample (10 μl) was fixed using 4% EM-grade PFA (10 μl) for 30 min at room temperature. Following fixation, a carbon-coated copper grid with type 200 mesh (Electron Microscopy Sciences, CF200-Cu-25) was allowed to adsorb to the EV/PFA mixture for 20 min. The grid was then placed on a drop of PBS to wash and then laid upon a drop of 1% glutaraldehyde in 1X PBS for 5 min. The grid was washed extensively (5-7 times x 2 min) on deionized water droplets, blotting gently in between each wash at a 45° angle on a piece of filter paper. The grid was then placed on a droplet of uranyl acetate replacement stain (Electron Microscopy Sciences, 22405) and let sit for 10 min. The grid was allowed to completely dry prior to imaging at 200 kV on a JEOL JEM 2100 LaB6 TEM.

### Gap Closure Assay

P4 HUVECs were seeded in gelatin-coated 96-well plates at 15,000 cells/well in EGM2 and allowed to grow until a uniform monolayer was formed (24 h). The cell monolayer was denuded using an AutoScratch (BioTek Instruments; Winooski, VT, USA). Cells were then washed once with 1X PBS and incubated with endothelial cell basal medium (EBM2; PromoCell C-22221) supplemented with 0.1% FBS for 2 h to serum starve the cells. After serum starvation, medium was replaced with fresh EGM2 (positive control), EBM2 (0.1% FBS) without EVs (negative control), or EBM2 with the addition of EVs at 5E9 EVs/mL based on NTA quantification or 200 μg/mL of EVs based on BCA quantification of surface protein. EBM2 or EGM2-treated cells were used as negative or positive controls, respectively. The closure of the cell gap was imaged at 0 h and 20 h using a Nikon Eclipse Ti2 Microscope at 2x magnification. Overall gap closure was determined as a percentage of area covered by HUVECs versus the gap area after 20 h using ImageJ as previously described [25].

### Tube Formation Assay

24-well plates were coated with 100 μl of growth factor reduced Matrigel (Corning, 354230) and incubated at 37°C for 30 min. P4 HUVECs were then seeded with EGM2 media (positive control), EBM2 media (0.1% FBS) devoid of EVs (negative control), or EBM2 media (0.1% FBS) with EVs (5E9 EVs/mL) on top of the Matrigel. At six hours, cells were imaged using a Nikon Eclipse Ti2 Microscope at 2x magnification. The number of fully-closed loops formed by the HUVECs were counted and recorded as a proxy for the formation of capillary-like structures in wound healing.

### EV Uptake via Flow Cytometry and Confocal Imaging

MSC EVs from flask and bioreactor culture were labeled separately with lipid membrane PKH67 dye (Sigma-Aldrich, PKH67GL) as previously described [23]. Briefly, EVs in PBS suspension were spun down at 8,000 × *g* in a 300 kDa MWCO filter and resuspended in 200 μl of Diluent C to wash away buffer-related salts. EVs were then spun down again at 8000 × *g* and resuspended in 250 μl of Diluent C per 200 μg of EVs. A mock dye treatment was prepared by mixing PBS devoid of EVs with diluent C. EVs and the mock dye solution were labeled at 1:1 ratio using 4 μM of PKH67 dye diluted in diluent C solution and incubated for 5 min with frequent mixing at room temperature every minute. 1% BSA prepared in diluent C was added to the EV or PBS/PKH67 dye mix at 1:1:1 ratio and incubated for another 1 min at room temperature to quench any leftover dye. Samples were then concentrated using protein concentrators (100 kDa MWCO; ThermoFisher Scientific, 88524) to 500 μl and centrifuged at 10,000 × *g* for 10 min at 4°C to remove any protein aggregates formed during concentration. To remove any non-EV associated dye aggregates, the samples were then run through size exclusion columns (Izon; qEV original 35 nm, ICO-35) per the manufacturer’s instructions. The first four fractions after the void volume were collected and used in further experiments (**Figure S1**). These fractions were concentrated (100 kDa MWCO), resuspended in 1X PBS, and sterile filtered (0.2 μm). Particle concentration was determined via NTA. Approximately 500,000 HUVECs in a 6-well plate were treated with 5E9 EVs/mL of labeled flask or bioreactor EVs or equal volume of mock dye solution in duplicates, while untreated HUVECs (PBS vehicle) were used as a negative control. HUVECs treated with the mock dye solution were used to assess uptake of any dye aggregates. After 24 h at 37°C, media were collected from each treatment to collect nonadherent cells. Adherent cells were detached using 1 mL of Accutase (ThermoFisher Scientific, A1110501), then added to the collected nonadherent cells, and spun down at 220 × *g* for 5 min. Cells were washed twice in 500 μl of FACS buffer (1% BSA in 1X PBS) and then resuspended in 250 μl of FACS buffer and strained through a cell strainer (40 μm) for further analysis. Samples were analyzed on an Amnis ImageStream X Mark II Imaging flow cytometer and analyzed using the IDEAS software. Live cell gating was done first in forward light scatter/side scatter, and PKH67 populations were gated from the live cell population.

In a concurrent experiment, HUVECs were seeded onto gelatin-coated coverslips (0.1%) in a 6-well plate. After 16 h, cells were treated with the PKH67 mock dye solution or PKH67-labeled flask or bioreactor EVs (5E9 EVs/mL). Cells treated with PBS devoid of dye or EVs acted as an additional control. After a 24 h incubation, HUVECs were fixed with 4% PFA for 15 min. Cells were washed three times with 1X PBS and then permeabilized for 5 min using a 0.5% Triton-X-100 solution. Following permeabilization, HUVECs were washed three times with 1X PBS and then blocked at room temperature for 1 h with a 2.5% goat serum (Abcam, ab7481) solution. Cellular actin was stained in a 1:100 dilution of Alexa Fluor Plus 647 Phalloidin (Invitrogen, A30107) coupled with a nuclear stain using a 1:10,000 dilution of DAPI (Cayman Chemical Company, 14285) for 30 min at room temperature. HUVECs were then washed three times in 1X PBS and imaged using an Olympus FLUOVIEW FV3000 confocal laser scanning microscope (Olympus, Shinjuku City, Tokyo, Japan) at 60x magnification.

**Figure S1.**
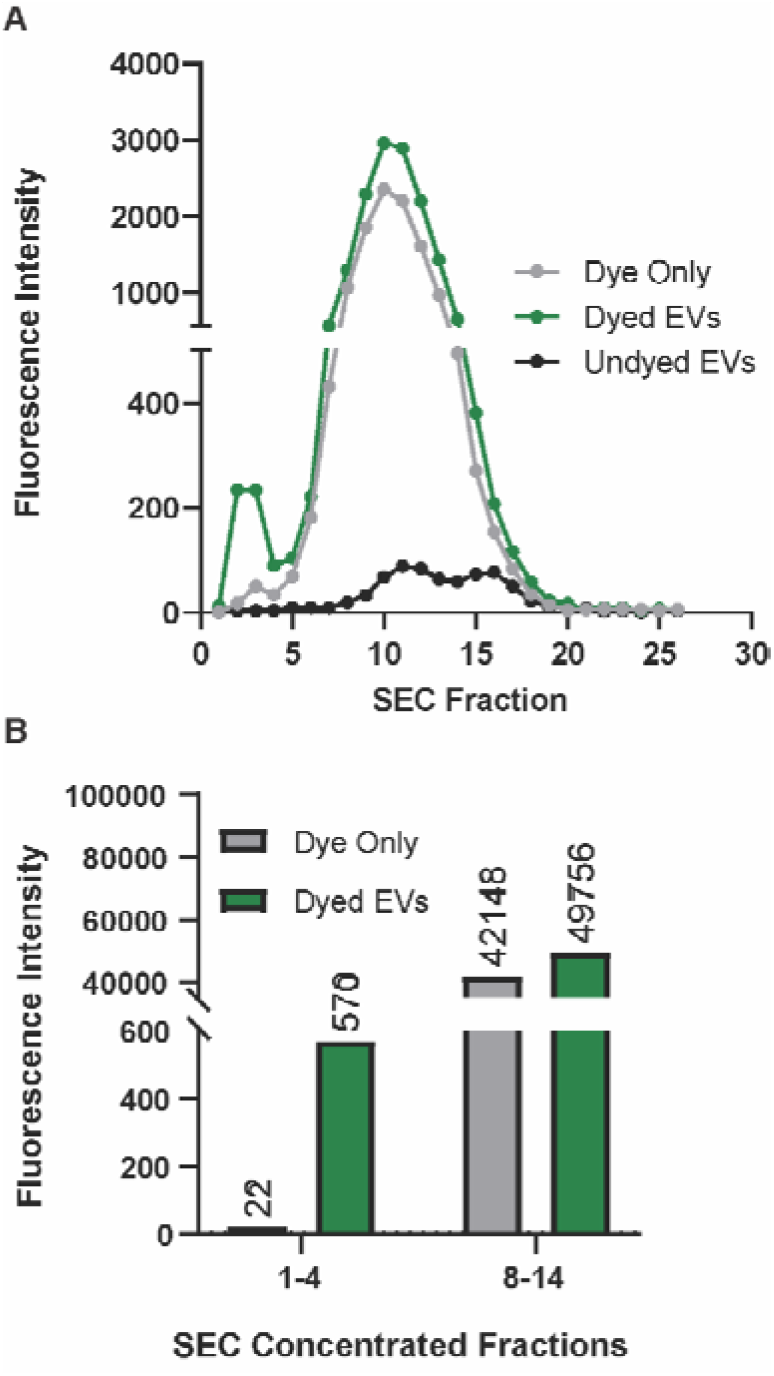
Size exclusion chromatography (SEC) can effectively remove PKH67 dye aggregates. (A) Fluorescence readings of each fraction from dye only (no EVs), dyed EVs, and undyed EVs (no PKH67) subjected to SEC. EVs elute in the first four fractions. (B) Fluorescence readings from the pooled fractions. Dye aggregates appear to elute in later fractions.

### MALDI-TOF Mass Spectrometry

Fresh MSC EVs from the flask and perfusion bioreactor (5 mL/min) conditions were washed in sterile deionized water to minimize the presence of buffer-related salts and were resuspended in a final volume of 300 μl of deionized water. BCA and NTA were performed and EVs were subjected to MALDI-TOF analysis on a MicroFlex LRF MALDI-TOF mass spectrometer (Bruker, Bremen, Germany). Each EV sample was mixed 1: 1 with sinapic acid matrix (Millipore Sigma, 85429; 20 mg/mL in 50% acetonitrile, 50% water, 0.1% trifluoracetic acid v/v/v) and 1 μl of the sample/matrix solution was deposited on a MicroFlex AnchorChip plate (Bruker, Bremen, Germany) in triplicate and dried in a vacuum. The analysis was performed under linear positive mode using the following parameters as previously described [26]: 70% laser intensity, laser attenuator with 35% offset and 40% range, accumulation of 500 laser shots, and 10.3 detector gain. Mass calibration was completed using cytochrome c (2 mg/mL; Sigma Aldrich, C8857) and myoglobin (2 mg/mL; Sigma Aldrich, M0630).

### mRNA Isolation and Profiling

P4 HDMECs were treated with PBS (control) or 200 μg/mL MSC EVs from flask or bioreactor (5mL/min) culture. After 24 h, HDMEC RNA was isolated using RNeasy kits (Qiagen, 74106) and cDNA was prepared using the iScript cDNA synthesis kit (Bio-Rad, 1708891). Following the manufacturer’s protocol, approximately 5 ng of RNA converted to cDNA was used per 10μL well reaction in the Wound Healing PCR Array (Bio-Rad, 10034601). Real-time PCR reaction was prepared using SsoAdvanced Universal SYBR Green Supermix (Bio-Rad, 1725271). Quantitative PCR was performed using an ABI 7900 Fast HT machine, with recommended thermal cycler settings for the SYBR Green Supermix. Data were analyzed using the ΔΔCt method. β-actin mRNA was used as a housekeeping control (ΔCt = Ct target gene – Ct β-actin) and results were normalized to PBS (ΔΔCt =ΔCt Bioreactor or Flask – ΔCt PBS). Data are shown in a heat map, where negative values indicate higher expression and positive values indicate lower expression compared to PBS treated group.

### *in vivo* Studies

All animal experimental protocols were approved by the Johns Hopkins University Animal Care and Use Committee, followed the Johns Hopkins University ACUC (Protocol RA18M86), and performed at Johns Hopkins University. A total of 24 db/db mice (40-50 g) from Jackson Laboratory (Bar Harbor, ME) were utilized (eight mice per treatment). All mice were anesthetized with 1.5% isofluorane (Baxter Healthcare Corporation, Deerfield, IL) and had their entire dorsum shaved where a singular 8 mm punch biopsy (Integra, Plainsboro, NJ) was performed. Buprenorphine (0.05 mg/kg) was given subcutaneously on days 0, 1, and 3 to help reduce any unnecessary discomfort. Treatments (i.e., flask EVs, perfusion bioreactor EVs, or PBS devoid of EVs) were injected four times around the wound in a cross pattern on day 3. Each dose contained 50 μg of EVs in a total volume of 50 μl or 50 μl of PBS without EVs. Photographs of the wounds were taken on days 0, 3, 7, 10, 12, and 14 and wound size was quantified by digital processing using Adobe Photoshop. On days 7, 10, 12, and 14, the wounds were debrided of the eschar to allow clear visualization. Wound size was calculated as the percentage of area of the wound versus the wound size on day 0.

### Histology

On day 21, healed tissues were biopsied using a 12 mm punch biopsy. The tissue was then cut down the center of the wound area and placed in a cryomold (Tissue Tek, 4557), where it was covered with OCT medium (Leica, 3801480). The tissue-containing cryomold was then placed onto a metal surface and cooled with dry ice and ethanol until the OCT medium solidified. The samples were stored at −80°C for less than 1 week prior to sectioning (10 μm thick) using a CM1950 Cryostat (Leica, Wetzlar, Germany). Tissue sections were fixed and permeabilized in 1:1 methanol:acetone solution at −20°C. Sections were then stained with H&E using a previously described protocol [8]. In brief, sections were washed for 2 min in deionized water, hematoxylin (VWR, 75810-352) for 3 min, deionized water for 1 min, differentiated in 4% HCl in 95% ethanol for 1 min, washed for 1 min in deionized water, bluing for 1 min in 1% NaCO3, washed for 1 min in deionized water, 95% ethanol for 1 min, eosin (VWR, 75810-354) for 45 s, 95% ethanol for 1 min, 100% ethanol for 1 min twice, and xylene for 2 min twice. Sub-X Mounting Medium (Leica, 3801740) was used to add a # 1.5 micro coverslip (VWR, 48393-195) on top of the tissue section and was sealed using clear fingernail polish. The tissue sections were then imaged using a Nikon Ti2 Microscope at 10x magnification.

In addition, 10 μm tissue sections were subjected to CD31 staining in order to detect newly formed blood vessels using a previously established protocol [8]. In short, slides were washed with tris-buffered saline (TBS) for 2 min, pre-blocked with 1% bovine serum albumin (Sigma Aldrich, A2058)/5% donkey serum (Sigma Aldrich, D9663) in TBS for 30 min, incubated with CD31 primary antibody (Abcam, 28364) at 1:50 in blocking solution for 60 min at room temperature, washed with TBS for 5 min twice, incubated with Alexa Flour 647 donkey anti-rabbit secondary antibody (Invitrogen, A31573) for 60 min at room temperature, and washed with TBS for 5 min twice. Coverslips were mounted over the tissue section as described in the previous section. Fluorescent images were taken using a Nikon Eclipse Ti2 Microscope at 10x magnification. The numbers of vessels were counted, and the area of tissue was quantified using ImageJ.

### Statistics

Data are presented as mean ± standard error of the mean (SEM). Each experiment involved three biological replicates, unless otherwise specified. One-way ANOVA with Tukey’s or Holm-Šídák’s multiple comparisons tests were used to determine statistical differences (p < 0.05) among groups in the EV characterization data, the *in vitro* tube formation assays, and *in vivo* vessel density data. Two-way ANOVA with Tukey’s or Holm-Šídák’s multiple comparisons tests were used to detect statistical differences (p < 0.05) among groups across dosing schemes in the *in vitro* gap closure assay, across donors within the different culture conditions, and among groups in the *in vivo* wound healing experiments over time. All statistical analyses were performed using Prism 9.1 (GraphPad Software, La Jolla, CA). Notation for significance in figures are as follows: ns – p > 0.05: #, *, or § - p < 0.05; §§ or ** - p < 0.01; *** - p < 0.001; ****, §§§§, ####, or †††† - p < 0.0001.

## Results

### Culture of MSCs in 3D-Printed Scaffold Perfusion Bioreactor

MSCs are physiologically subjected to interstitial flow that exerts low levels of shear stress that has been shown to impact their growth kinetics as well as differentiation [27, 28]. We designed and 3D-printed scaffolds with a total surface area of 50 cm^2^ consisting of a pillared array (**Figure 1A**) that permitted medium and gas flow through gas permeable tubing (**Figure 1C**). Computational fluid modeling was carried out for 1, 5, and 10 mL/min to evaluate shear stress (**Figure 1D**) and flow trajectories (**Figure 1E**) throughout the scaffold. Shear stress and flow trajectory heat maps show the distribution of shear stresses across the scaffold growth surfaces, with highest values found closed to the entrance and exit of the scaffold. Average shear stress values for 1, 5, and 10 mL/min were calculated to be 1.6×10^-4^, 3×10^-3^, and 1.2×10^-2^ dyn/cm^2^, respectively, all of which are below physiological interstitial shear stress levels of 0.01Pa (0.1 dyn/cm^2^) [29]. For cell viability analysis, MSCs were fixed and stained with AlexaFluor 488 Phalloidin after exposure to dynamic culture at 0, 1, 5, or 10 mL/min. Images confirmed the presence of live cells throughout the scaffold base and along the side walls of the pillars (**Figure 1F**) that were subjected to no flow or flow rates of 1 and 5 mL/min. At a flow rate of 10 mL/min flow rate, very few cells remained adhered to the scaffold.

**Figure 1.**
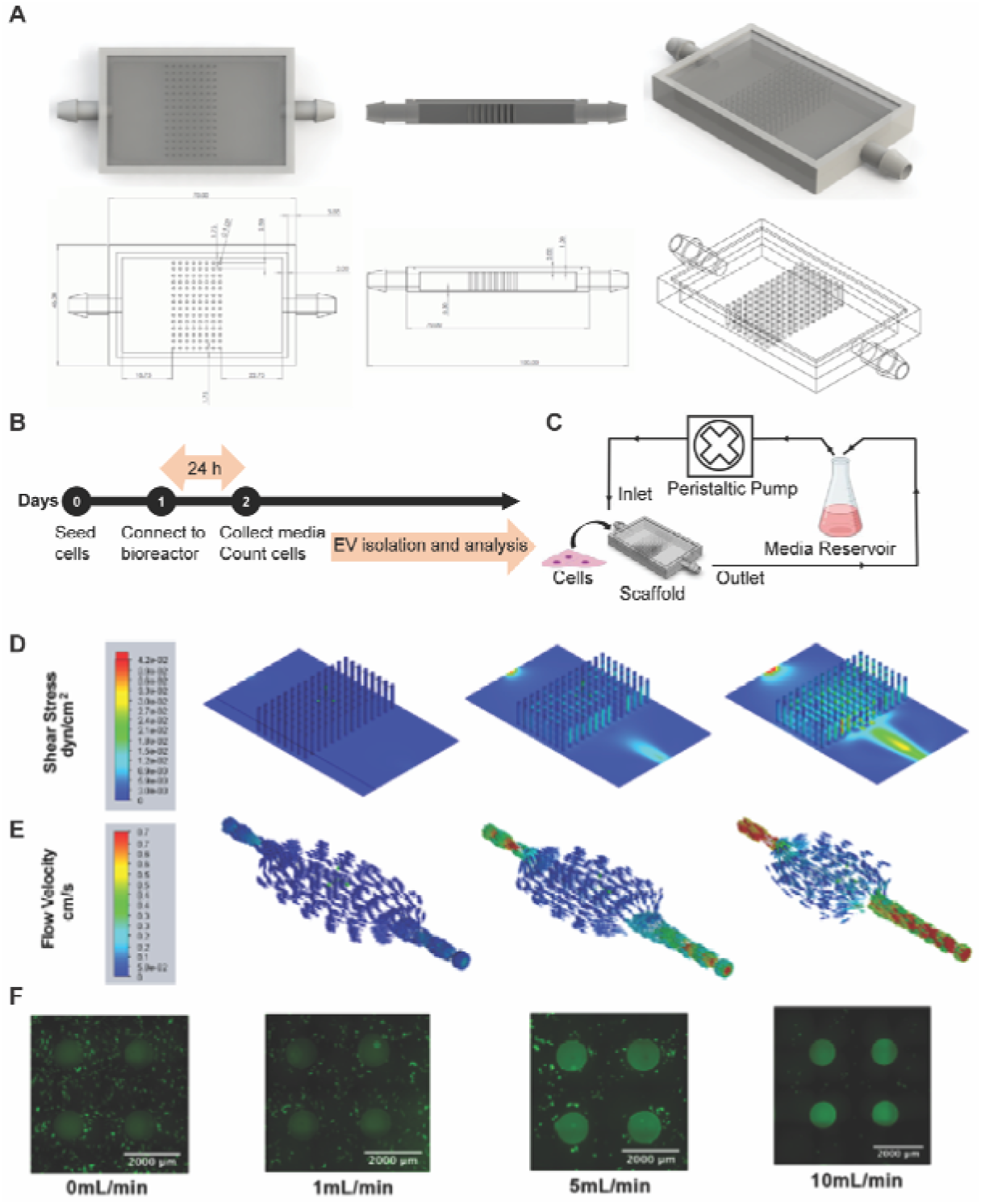
Perfusion bioreactor setup and scaffold characterization. (A) Cross-sectioned image of the scaffold generated in SolidWorks showing dimensions of the scaffold as well as individual pillar height, diameter, and spacing. (B) Timeline of the perfusion bioreactor experimental procedure. (C) Schematic representation of the perfusion bioreactor setup. A scaffold seeded with cells is connected to a peristaltic pump and media reservoir using gas-permeable tubing. Inlet and outlet of the media circulates at 1, 5, or 10 mL/min flow rate. (D) Visual representation of varying levels of shear stress and (E) flow trajectories at different areas within the scaffold as evaluated by computational flow simulation are shown for 1, 5, and 10 mL/min. (F) Images of the phalloidin-stained scaffold seeded with MSCs after 24 h of dynamic culture.

### MSC EV Production is Enhanced by Perfusion Bioreactor Culture

We first aimed to assess the impact of increasing flow rates on MSC EV production. EVs were isolated from MSCs cultured in flask, scaffold (0 mL/min), or perfusion bioreactor conditions (1, 5, or 10 mL/min) for 24 h. The flow rates were selected based on CFD analysis to determine maximum shear stress to not exceed physiological interstitial levels of 0.1 dyn/cm^2^, which has been shown to induce MSC differentiation into osteogenic phenotype [28]. NTA revealed no significant difference in EV size distribution. Peak size values for EVs from all culture conditions were approximately 160 nm (**Figure 2A**). More than 90% of the total EV populations from all culture conditions were within the typical exosomal diameter range (40-200nm) (**Figure 2A**) [30]. The highest level of EV production per cell occurred when MSCs were exposed to a 5 mL/min flow rate (9.8E5 ±9E4), which was 83-fold, 28-fold, 3-fold, and 2.5-fold higher than flask, 0, 1, and 10 mL/min, respectively (flask: 1.18E4 ± 9.3E3; 0 mL/min: 3.5E4 ± 5.2E3; 1 mL/min: 3.0E5 ± 2.7E4; 10 mL/min: 3.9E5 ± 4.6E4) (**Figure 2B**). This trend of elevated EV production as determined by NTA was retained across various tissue origins (i.e., bone-marrow and adipose-derived MSCs) as well as different donors (**Figure S2**). EV production per cell was also evaluated using a CD63-specific ExoELISA, which confirmed the significant increase in EV output in the bioreactor culture system at 1, 5, and 10 mL/min compared to flask culture (25-fold, 43-fold, and 47-fold) and 0 mL/min (23-fold, 39-fold, and 43-fold), respectively (**Figure 2C**). Interestingly, exoELISA results showed higher CD63^+^ EV production per cell at 10 mL/min compared to the total EV population per cell assessed by NTA (**Figure 2C**). As measured by BCA, average protein content per EV decreased 2-fold in the 5 mL/min bioreactor compared to the flask culture, but was not significant (p > 0.05) (**Figure 2D**). Protein content per EV was inversely proportional to EV production rates (**Figure 2D**).

**Figure 2.**
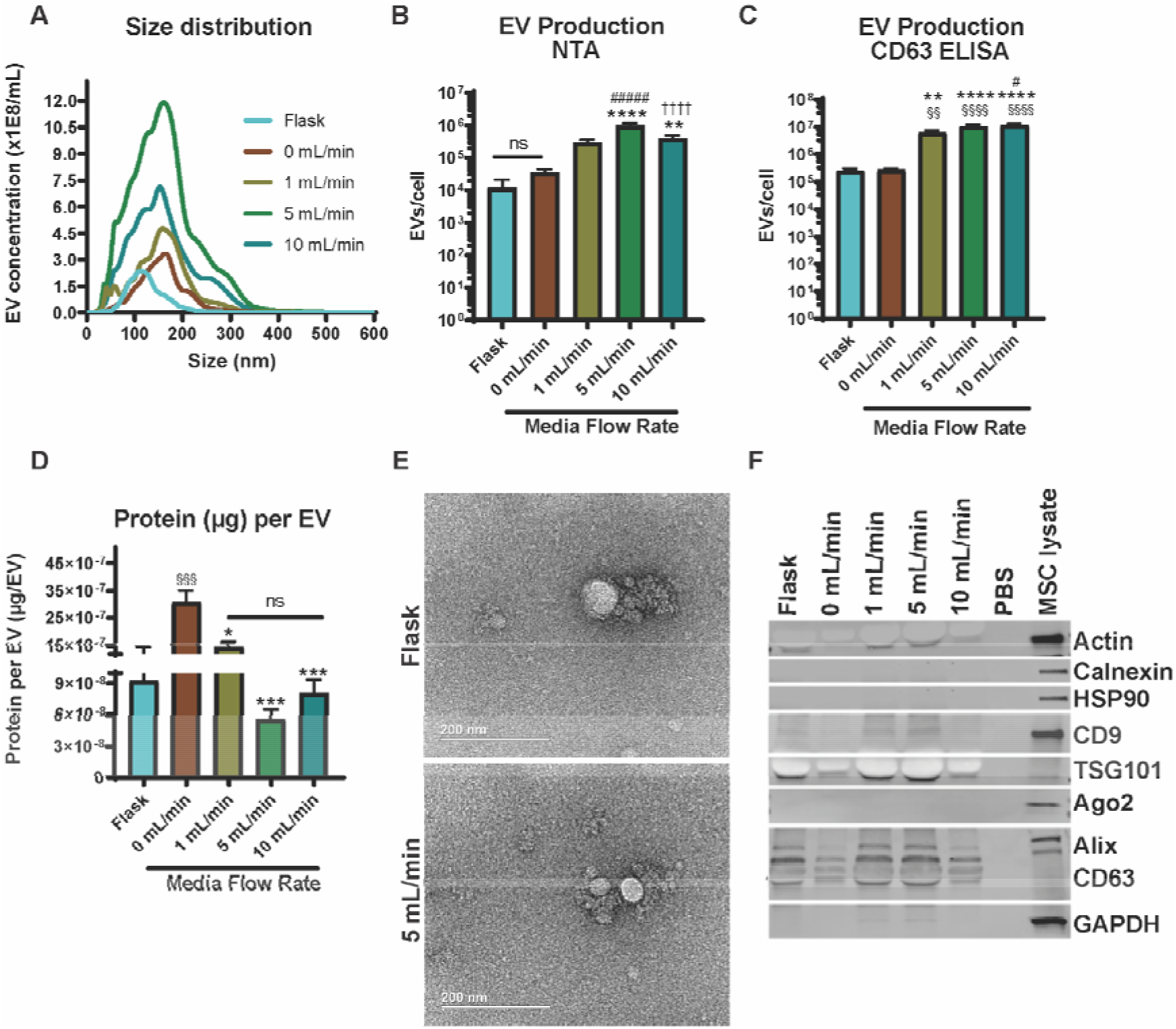
MSC EV production can be maximized in the bioreactor with a 5 mL/min flow rate. (A) Size distribution of MSC-derived EVs produced in the various culture conditions. Number of EVs produced per cell as measured using (B) nanoparticle tracking analysis (NTA) and (C) a CD63-specific ExoELISA. (D) The ratio of total surface protein amount (μg) to total number of EVs are shown. A significant increase in EV production (§ - compared to flask, * - compared to 0 mL/min, # - compared to 1 mL/min, † - compared to 5 mL/min) was recorded, but a decrease in protein content (p < 0.001) was observed for the 5 mL/min condition when compared with the 0 mL/min condition. (E) TEM images of EVs from MSCs either cultured in the flask or 5 mL/min conditions. (F) Immunoblots of MSC EVs from the various culture conditions for EV markers (CD63, Alix, TSG101, CD9), cell markers (Calnexin, HSP90, and Ago2), and control markers (Actin and GAPDH). Data representative of at least three independent experiments (N = 3). Statistical significance was calculated using a one-way ANOVA using Tukey’s multiple comparison tests (ns – p > 0.05: #, *, or § - p < 0.05; §§ or ** - p < 0.01; *** - p < 0.001; ****, §§§§, ####, or †††† - p < 0.0001).

**Figure S2.**
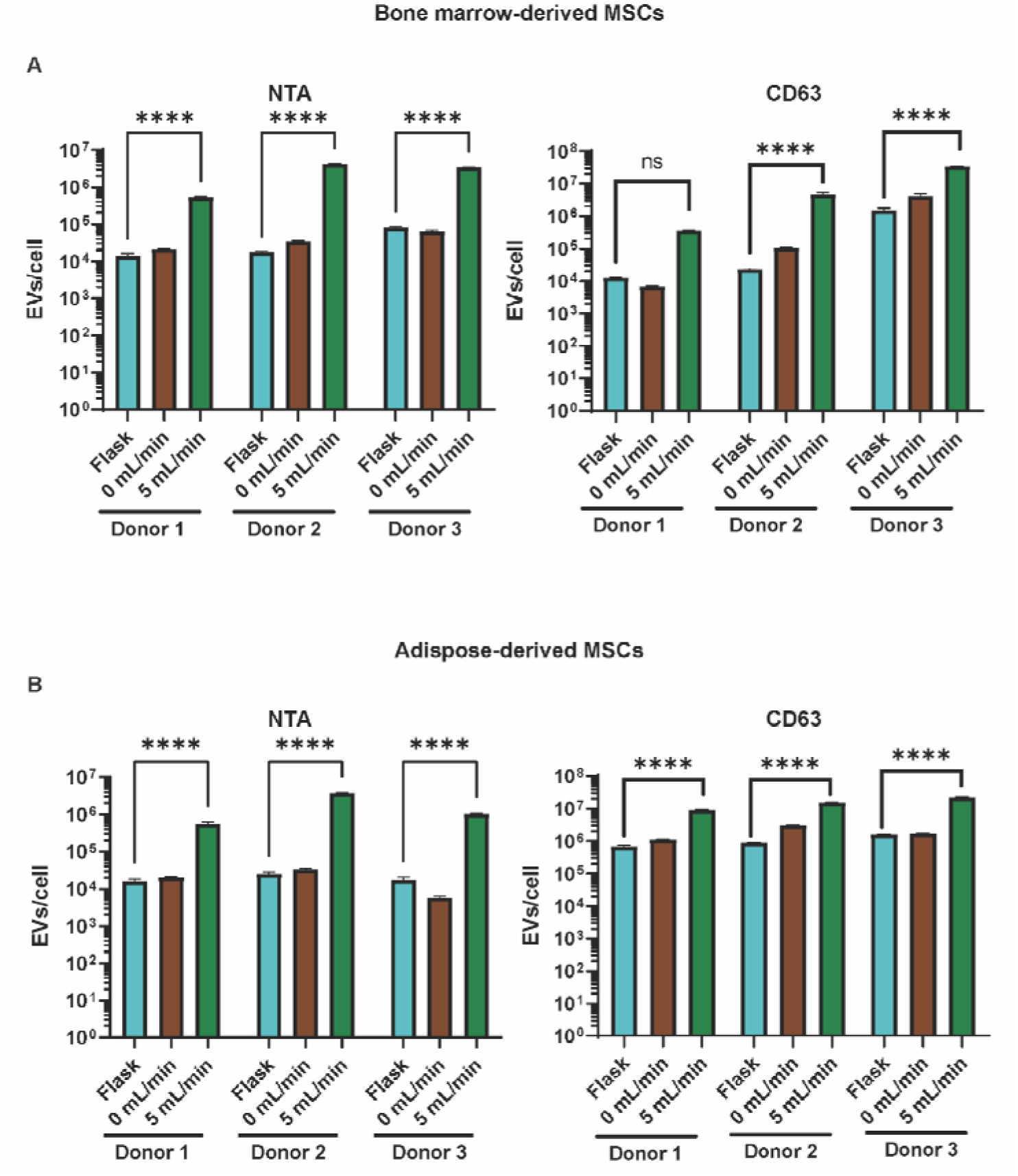
Flow-induced increase in MSC EV production is maintained across donors and tissue of origin. EV production from three donors of (A) bone marrow-derived MSCs and three donors of (B) adipose-derived MSCs as measured by NTA and a CD63 ELISA. A significant increase in EV production occurred when cells were cultured in the 5 mL/min bioreactor when compared with those maintained in flasks regardless of donor or tissue origin. Data representative of three independent experiments (N = 3). Statistical significance was calculated using a two-way ANOVA using Tukey’s multiple comparison tests (ns – p > 0.05: * - p < 0.05; ** - p < 0.01; *** - p < 0.001; ****- p < 0.0001).

We observed a 2-fold, 5-fold, and 4-fold decline in protein content per EV compared to 0 mL/min in the 1, 5, and 10 mL/min cultures, respectively. TEM images revealed no observable changes in MSC EV morphology when MSCs were cultured within the bioreactor (i.e., 5 mL/min) (**Figure 2E**). Immunoblot analyses confirmed the presence of EV markers (Alix, TSG101, CD63, and CD9) and absence of cellular debris markers (Calnexin, HSP90, and Ago2) in MSC EV samples for flask, and 0, 1, 5, and 10mL/min culture conditions (**Figure 2F**). The 5 mL/min flow rate was determined to be optimal as it was able to significantly increase EV production without dislodging cells (**Figures 1F and 2B**), and was thus used for further experiments.

### Enhanced EV Production in Perfusion Bioreactor Culture is Not Specific to MSCs

In an attempt to understand whether the observed boost in MSC EV production is the result of a cell-driven mechanoresponse, we cultured HEK293 cells within the system as they lack the abundance of mechanoreceptors inherently present in MSCs [31–33]. Interestingly, the HEK cells demonstrated a similar and significant uptick in EV production as well as a decreasing trend in protein per EV as flow rate increased (**Figure 3A, B**). HEK EV morphology was not altered by the culture conditions (**Figure 3C**). And immunoblotting confirmed the presence of EV markers (CD63, Alix, TSG101) and the absence of the cellular debris marker calnexin (**Figure 3D**).

**Figure 3.**
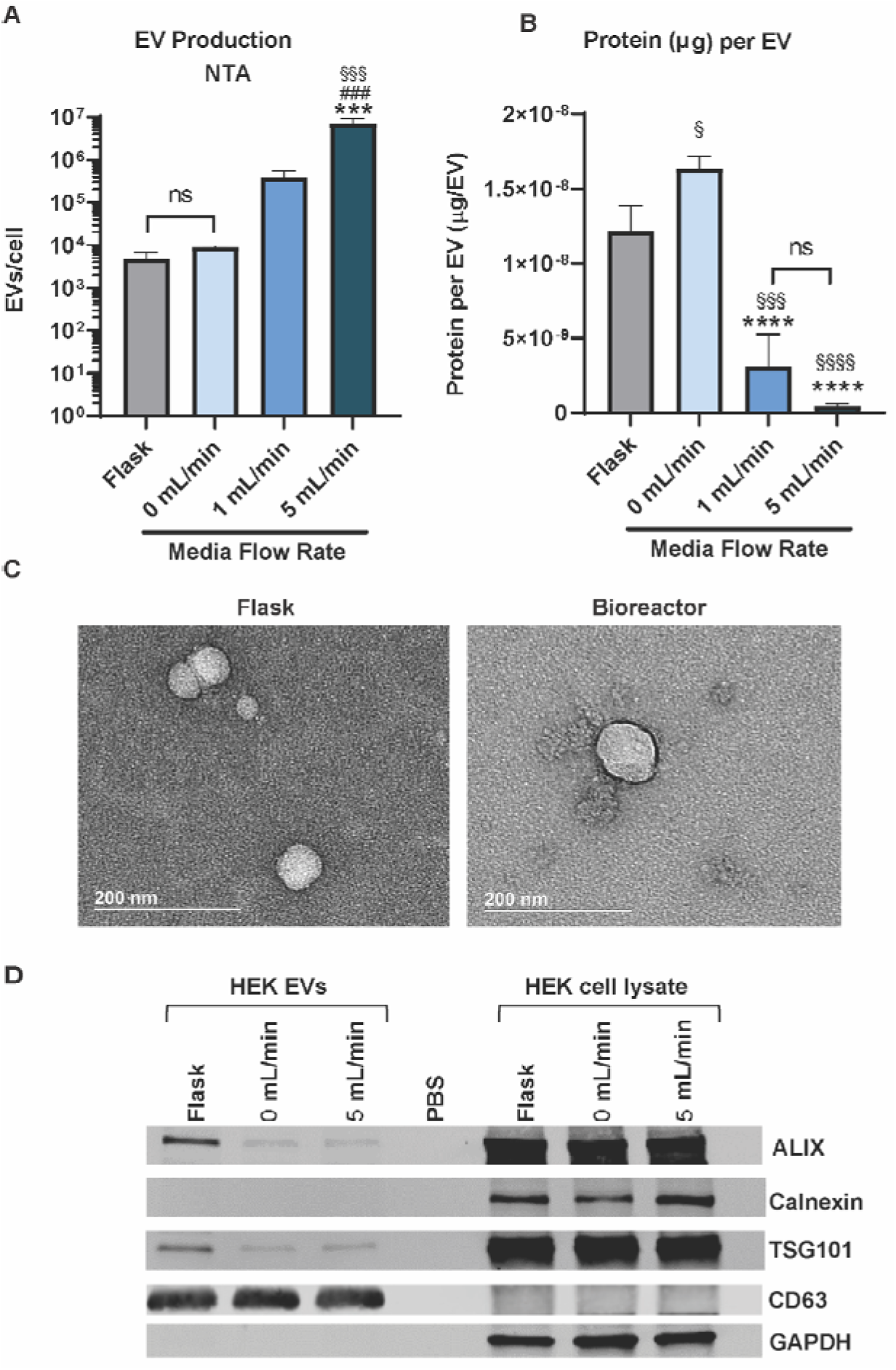
HEK293 EV production increases when cultured in perfusion bioreactor. (A) Size distribution of HEK-derived EVs produced in the various culture conditions. (B) Number of EVs produced per cell as measured using nanoparticle tracking analysis (NTA). (C) The ratio of total surface protein amount (μg) to total number of EVs are shown. A significant increase in EV production (§ - compared to flask, * - compared to 0 mL/min, # - compared to 1 mL/min) was recorded, and a decrease in protein content (p < 0.0001) was observed for the 5 mL/min condition when compared with the 0 mL/min condition. (C) TEM images of EVs from HEK cells maintained in the flask or bioreactor conditions. (F) Immunoblots of HEK EVs from the various culture conditions for EV markers (CD63, Alix, TSG101) and cell markers (Calnexin, GAPDH). Data representative of at least three independent experiments (N = 3). Statistical significance was calculated using a one-way ANOVA using Tukey’s multiple comparison tests (ns – p > 0.05: #, *, or § - p < 0.05; ***, §§§, or ### - p < 0.001; **** or §§§§- p < 0.0001).

### MSC EV *in vitro* Angiogenic Bioactivity is Maintained Following Perfusion Bioreactor Culture

Bioactivity of MSC EVs isolated from the flask and optimal perfusion bioreactor condition (5 mL/min) was subsequently assessed using an *in vitro* gap closure assay and a tube formation assay. In the gap closure assay, when treated with flask-derived MSC EVs based on EV count (i.e., 5E9 EVs/mL), HUVECs had significantly higher gap closure when compared with the negative control that was treated with basal media only (p < 0.01) (basal media: 22.23 ± 3.659; flask at 5E9 EVs/mL: 48.49 ± 1.737) (**Figure 4A, B**). Using the same dosing scheme, bioreactor-derived MSC EVs also significantly improved gap closure in comparison with the negative control (p < 0.0001) and the flask-derived EVs (p < 0.05) (bioreactor at 5E9 EVs/mL: 65.26 ± 5.926). When dosed at 200 μg/mL, MSC EVs derived from both the flask and bioreactor conditions significantly augmented gap closure in HUVECs when compared with the negative control (p <0.01 and p < 0.0001; respectively) (flask at 200 μg/mL: 42.31 ± 2.746; bioreactor at 200 μg/mL: 52.72 ± 1.318). This trend of maintained EV bioactivity was retained across donors of bone marrow-derived MSCs, whereas we observed no angiogenic activity from adipose-derived MSC EVs (**Figure S3**). Importantly, there was no significant difference between the dosing schemes in the respective culture conditions. Therefore, in all subsequent experiments, MSC EVs were dosed at 5E9 EVs/mL.

**Figure 4.**
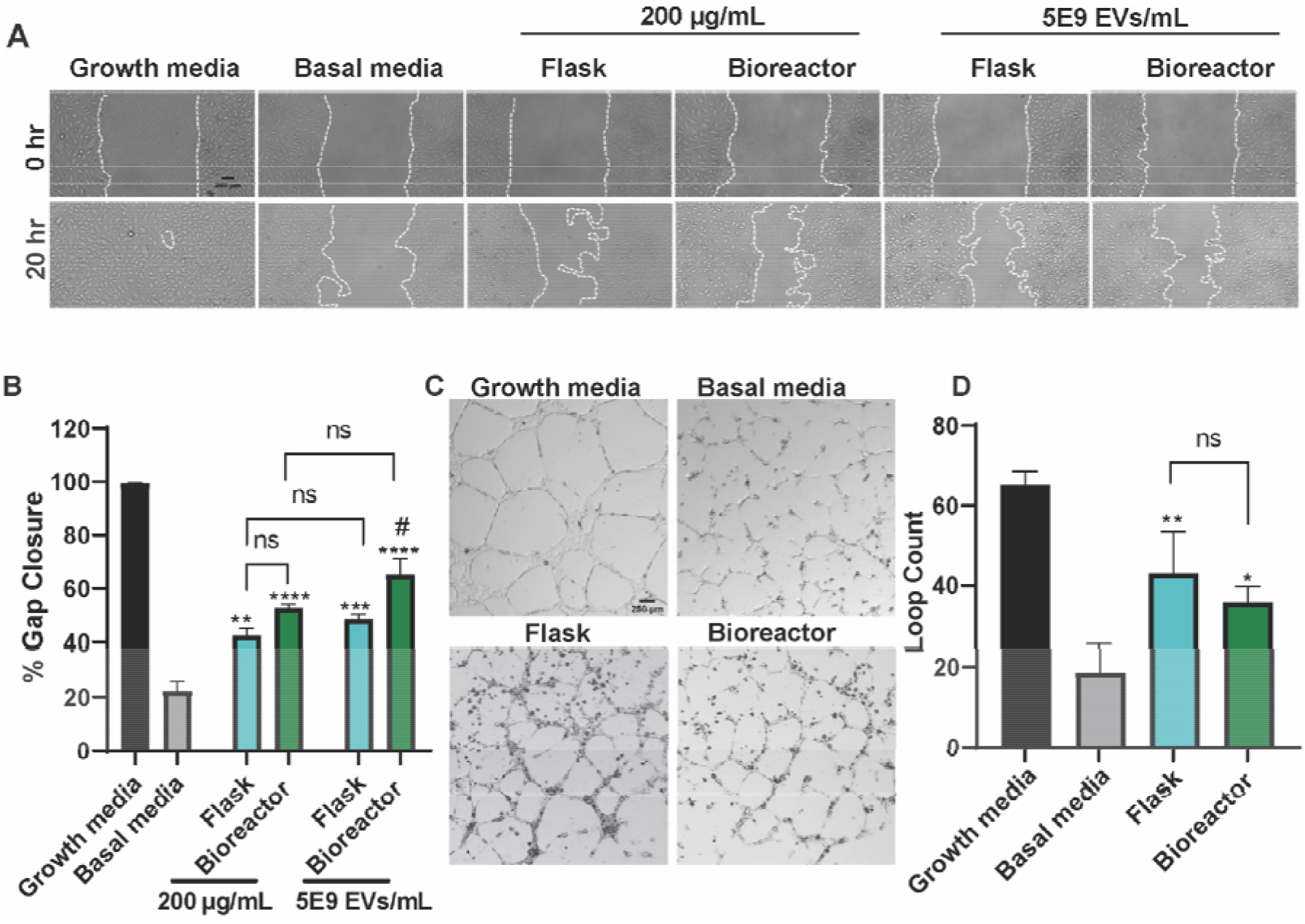
Perfusion Bioreactor culture maintains *in vitro* pro-angiogenic activity of MSC EVs. (A) Representative images of HUVECs in the gap closure assay at 0 and 20 h after treatment with growth media (positive control), basal media (negative control), or 5E9 EVs/mL or 200 μg/mL EVs isolated from MSCs in the flask or bioreactor (5 mL/min) culture conditions. (B) Cell gap area after 20 h as a percentage of gap area at 0 h. A significant increase in gap closure was observed when cells were treated with either flask or bioreactor EVs at 5E9 EVs/mL or 200 μg/mL (* - compared to basal, # - compared to flask). There were no significant differences detected between dosing schemes. (C) Representative images HUVECs in the tube formation assay 6 h after application of growth media (positive control), basal media (negative control), or 5E9 EVs/mL EVs from MSCs maintained in flasks or the bioreactor. (D) Results of the tube formation were quantified as the number of complete loops formed by HUVECs. Both flask and bioreactor EVs significantly increased the number of loops with no differences between flask and bioreactor EVs (* - compared to basal). All images and data are representative of three independent experiments (N = 3). Statistical significance was calculated using either a two-way (B) or one-way ANOVA (D) with Tukey’s multiple comparisons test (ns – p > 0.05; * or # - p < 0.05; ** - p < 0.01; *** - p < 0.001; ****- p < 0.0001).

In the tube formation assay, both flask- and bioreactor-derived MSC EVs significantly enhanced the number of loops formed by the HUVECs when compared with the negative control (p < 0.01 and p < 0.05, respectively; basal: 18.67 ± 3.93; flask: 43.00 ± 6.110; bioreactor: 35.67 ± 2.300) (**Figure 4C, D**). There was no significant difference between the flask and bioreactor EVs.

**Figure S3.**
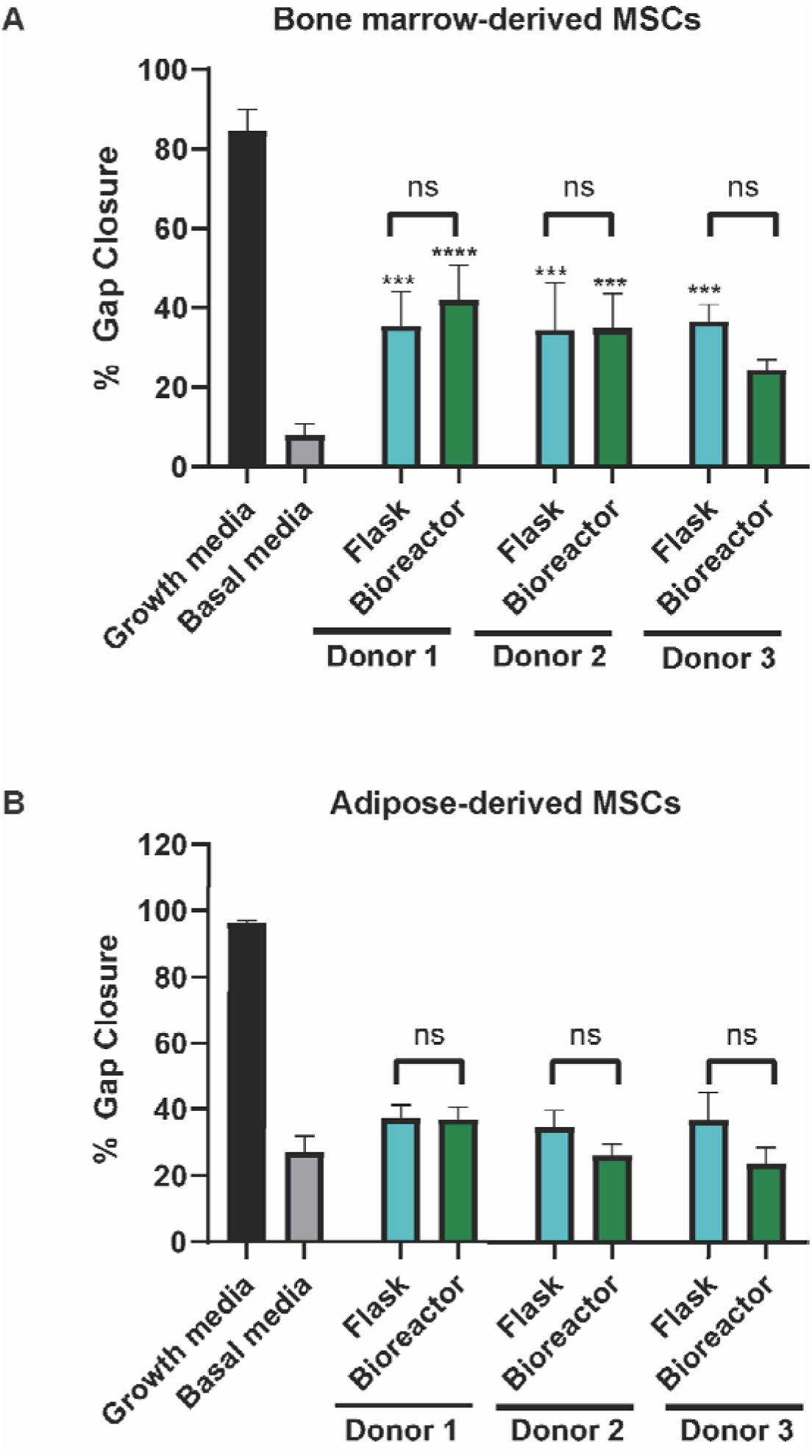
Angiogenic bioactivity is maintained across donors of bone marrow-derived MSCs. Cell gap area after 20 h as a percentage of gap area at 0 h. (A) A significant increase in gap closure was observed when HUVECs were treated with 200 μg/mL of EVs from three donors of bone marrow-derived MSC when compared with basal media (negative control). (B) EVs from adipose-derived MSCs had no effect on gap closure regardless of culture conditions. There were no significant differences between flask- or bioreactor-derived EVs from either tissue source. Data representative of three independent experiments (N = 3). Statistical significance was calculated using a two-way ANOVA using Tukey’s multiple comparison tests (ns – p > 0.05; *** - p < 0.001; ****- p < 0.0001).

### *In vivo* MSC EV Wound Healing Bioactivity is Enhanced via Perfusion Bioreactor Culture

The wound healing bioactivity of EVs from flask and bioreactor culture was assessed *in vivo* using a diabetic mouse wound healing model. An 8 mm punch biopsy excisional wound was created on the dorsum of each mouse. Three days post-injury, mice were injected four times around the wound with 50 μg of flask MSC EVs, bioreactor MSC EVs, or a PBS vehicle control. EVs isolated from MSCs cultured in the bioreactor generated a significant improvement in healing overall when compared with the vehicle control (p < 0.05) (**Figure 5A, B**). CD31 immunohistochemistry revealed that mice treated with bioreactor MSC EVs had significantly more CD31+ vessel structures when compared with animals treated with flask MSC EVs or PBS (p < 0.05; PBS: 4.7 ± 0.984; flask: 5.876 ± 1.636; bioreactor: 12.02 ± 2.423). There was no significant difference between the vehicle control and flask MSC EVs (**Figure 5C, D**).

**Figure 5.**
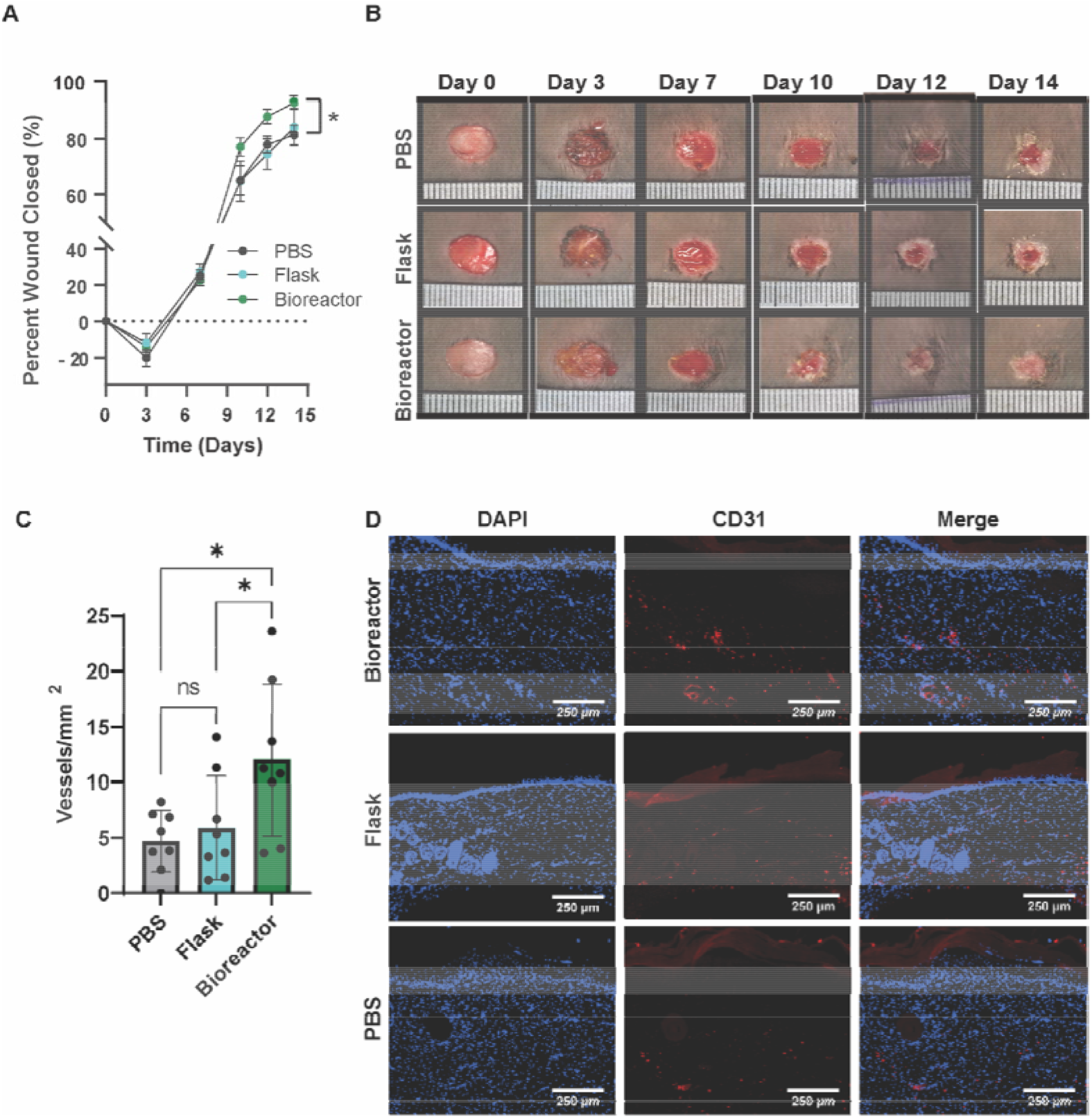
Perfusion bioreactor culture improves healing in a diabetic mouse excisional wound healing model. (A) Closure evaluated for wounds treated with flask MSC EVs, bioreactor MSC EVs, or PBS. Bioreactor MSC EVs improved overall wound healing compared with the PBS control (p < 0.05). (B) Representative images of mice wounds over the length of the experiment. (C) Number of CD31+ vessels in healed tissues isolated from mice on day 21 with (D) representative immunohistochemistry images. A significant enhancement of CD31+ vessels was apparent in mice treated with bioreactor MSC EVs (p < 0.05). Statistical significance was calculated using a (A) two-way ANOVA or (C) one-way ANOVA with Holm-Šídák’s multiple comparisons test (*p < 0.05; n = 8).

### Assessment of Perfusion Bioreactor Culture-Induced Enhancement of *in vivo* MSC EV Bioactivity

To determine whether any differences in MSC EV bioactivity observed *in vivo* could be attributed to disparate internalization into recipient cells, we evaluated EV uptake. HUVECs were treated with PKH67-labeled MSC EVs from flask or bioreactor culture for 24 h (**Figure 6A**). There was no difference in uptake of the EVs from the bioreactor culture (96.7%) compared to flask culture (98.6%) (**Figure 6B**), which was visually confirmed with confocal images of the recipient cells (**Figure 6C**). As flow has been shown to alter protein corona composition of nanoparticles which can in turn affect uptake [34, 35], we assessed the EV surface proteins using MALDI-TOF. However, the resulting spectra were highly reproducible over time (**Figure S4**) and showed no apparent differences between the flask and bioreactor MSC EVs (**Figure 7C**).

**Figure 6.**
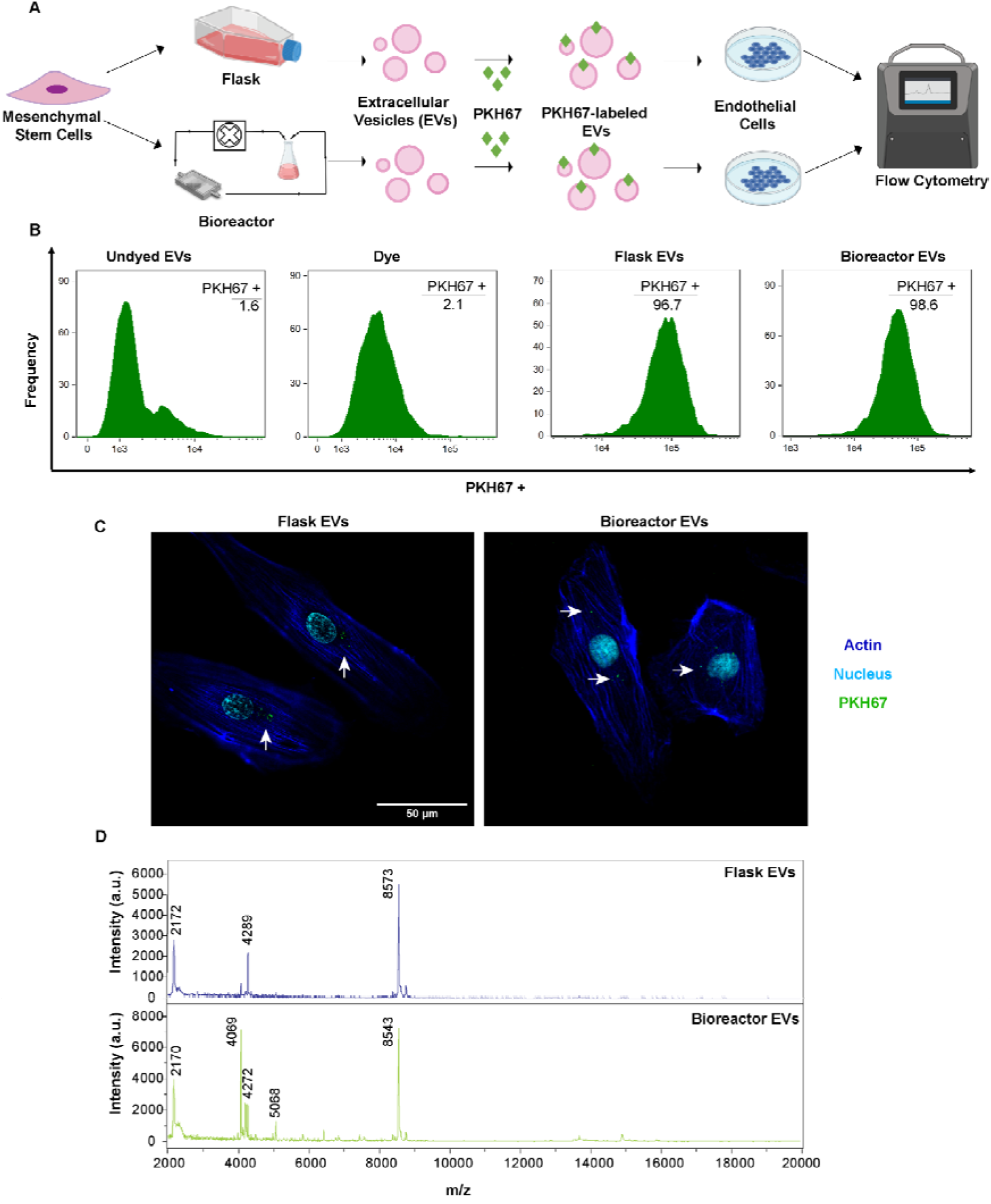
Assessment of HUVEC uptake and protein profile of flask and bioreactor MSC EVs. To assess EV uptake, flow cytometry analysis of PKH67-labeled EVs from MSCs cultured in flask vs. bioreactor conditions was conducted as shown in (A). (B) Histograms of frequency vs fluorescence intensity are shown for HUVECs that were treated with undyed EVs (no PKH67), just dye (no EVs), dyed flask EVs, or dyed bioreactor EVs. (C) Representative images of HUVECs after treatment with labeled EVs. White arrows denote the location of PKH67-labeled EVs. (D) MALDI-TOF mass spectra of flask and bioreactor-derived EVs. Data represent at least 2 independent experiments (N = 2).

**Figure 7.**
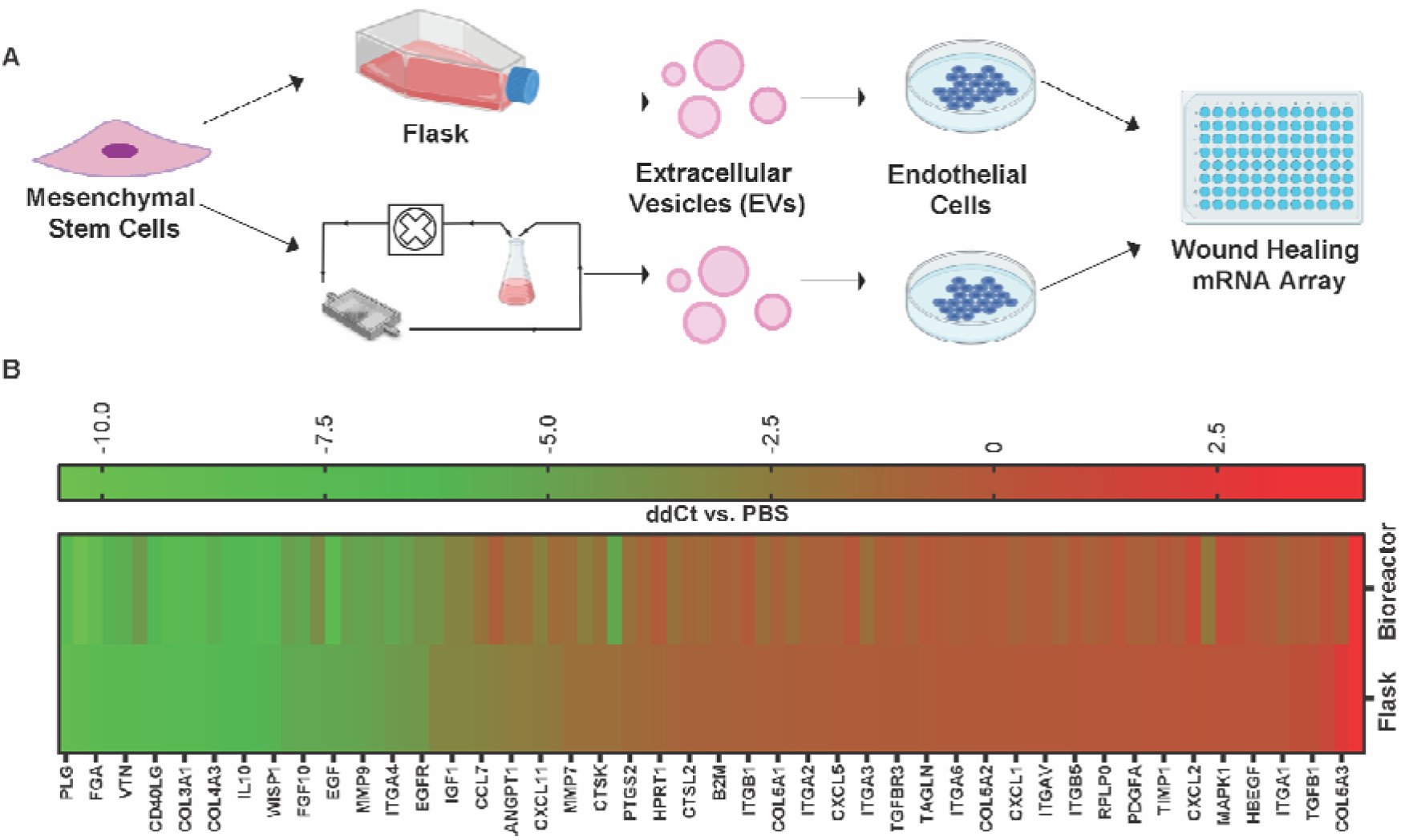
RNA profiling of HDMECs treated with MSC EVs from flask vs perfusion bioreactor culture. (A) Experimental schematic of wound healing specific gene expression in HDMECs treated with MSC EVs from flask or bioreactor culture. (B) Heat map representing gene regulation profile of 88 genes associated with wound healing. Data shown as ΔΔCt = (ΔCt Flask or Bioreactor – ΔCt PBS), where ΔCt = (Ct target gene – Ct β-actin). No significant difference between RNA profiles of HDMECs treated with flask or bioreactor derived MSC EVs was observed (data not shown). Significance was tested using one-way ANOVA with Bonferroni’s multiple comparison test. Data are representative of two independent experiments (N = 2).

Next, to assess any variance in RNA profiles of recipient HDMECs due to flask vs. bioreactor MSC EVs, we analyzed expression of 88 genes associated with wound healing (**Figure 7A**). Among the 88 genes, 34 were upregulated and only 2 were downregulated more than 2-fold compared to PBS for both flask and bioreactor EV groups. Between flask and bioreactor EV groups, 3 were upregulated (EGF, FGF, and COL5A3) and 3 were downregulated (VTN, ITGB6, and IL2) in the bioreactor EV group by more than 2-fold, though none showed significance (p > 0.05) (**Figure 7B**).

**Figure S4.**
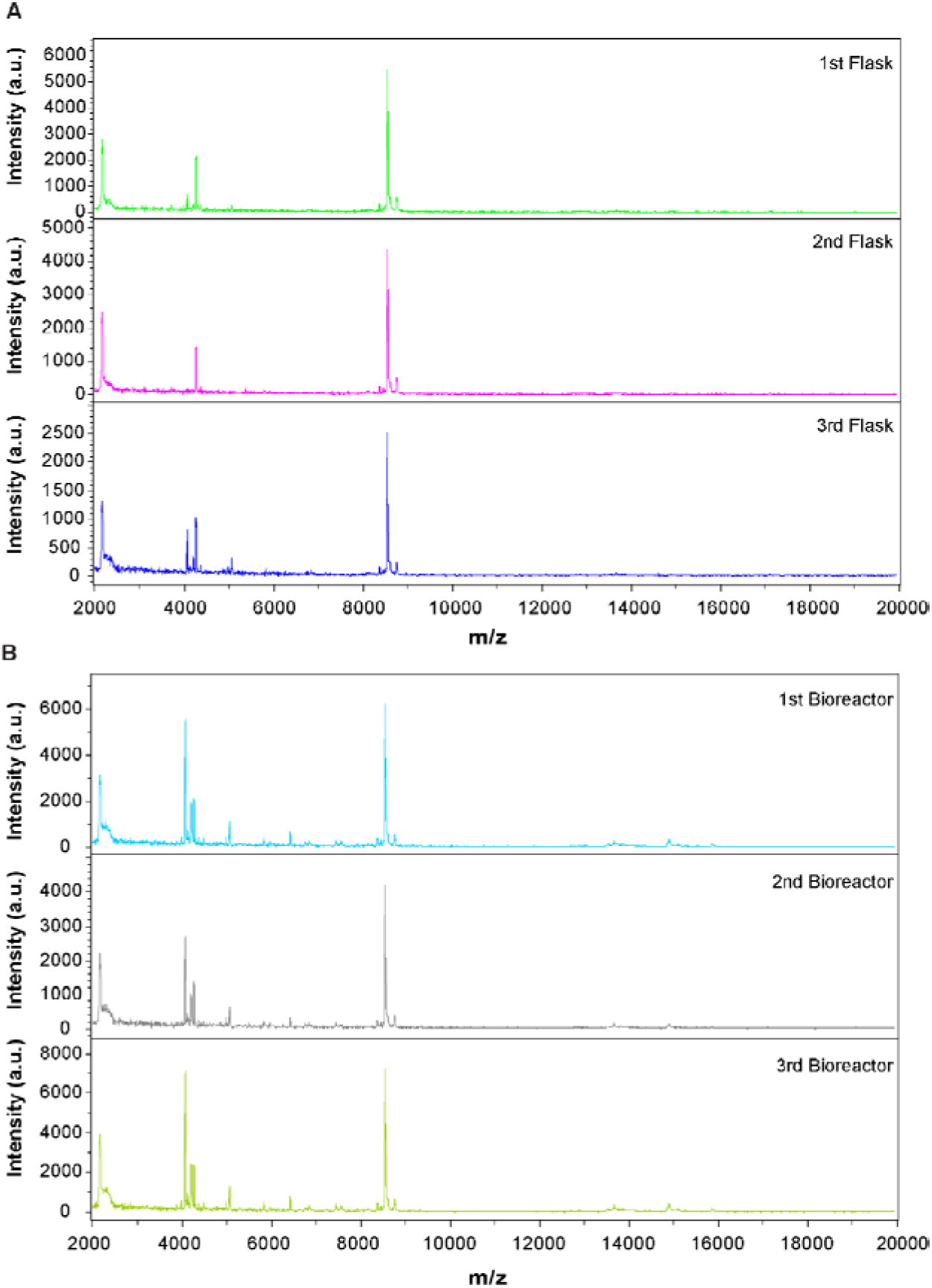
MALDI-TOF mass spectra of three separately prepared batches of flask and bioreactor EVs.

## Discussion

The lack of a scalable manufacturing platform has remained a substantial hindrance to the clinical translation of EV-based therapies [11, 36, 37]. The most commonly employed solution to this problem is the use of hollow-fiber technology, which is highly effective for optimizing yield [11, 38], but has limited versatility with regard to control of the various biophysical cues that can most influence EV properties, such as shear stress and substrate architecture. Engineering advancements, such as 3D printing, can allow for a more reductionist and tunable approach that enables control over these cues in a manner amenable to large-scale production. In a previous study, we leveraged 3D-printing to create a scaffold-perfusion bioreactor that significantly increased the secretion of EVs from human dermal microvascular endothelial cells (HDMECs) [18]. Inspired by these results, we attempted to use the perfusion bioreactor to enhance the therapeutic efficacy of MSC EVs, as MSCs are another mechanosensitive cell type with many therapeutic applications [20, 39]. In this study, the scaffold was redesigned to incorporate a pillar configuration previously implemented to create a highly-adaptable and physiologically-relevant microenvironment specifically for human MSCs [40]. In addition, when perfused with media, this design allowed for the use of flow rates that produced shear stress values well below levels previously reported for the *in vivo* stem cell niche (i.e., 0.1 – 1 dyn/cm^2^) and well below values known to induce MSC differentiation [27, 29, 41] (**Figure 1D**).

While implementing the bioreactor system, we found that a flow rate of 5 mL/min (3×10^-3^ dyn/cm^2^) appeared optimal as it significantly increased MSC EV production by 83-fold when compared with flask culture and by 2.5-fold when compared with a flow rate of 10 mL/min (1.2×10^-2^ dyn/cm^2^) (**Figure 2B**) without causing the significant cell detachment that was observed in the 10 mL/min condition (**Figure 1F**). However, when an ELISA was used to examine the levels of tetraspanin CD63, which is considered an indicator of EVs originating from the multi-vesicular body (i.e., exosomes) [42], the 5 mL/min produced only a 43-fold increase in CD63^+^ EVs while the 10 mL/min condition produced a very similar 47-fold increase in CD63^+^ EVs when compared with flask culture (**Figure 2C**). We also show a significant reduction in total protein content per EV as flow rate is increased from 0 to 5 mL/min, which seemed to saturate at 10 mL/min (**Figure 2D**). Protein per EV is often used as a proxy for EV purity, with less protein per EV being considered a gauge of higher purity [43]. We have previously reported similar phenomena in which HDMECs within a perfusion bioreactor produced a significant yet smaller increase in CD63^+^ EVs in comparison with total EVs along with a purer EV population [18]. Altogether, these results indicate that the bioreactor favors the secretion of other EV sub-populations such as microvesicles, which are known to originate from the cell membrane and may be composed of a relatively lower amount of CD63 [44, 45]. Indeed, shear stress is known to increase cell membrane tension which results in augmented exocytotic activity [46, 47]. Overall, the increased MSC EV production we observed mirrors the results of previous studies by other groups that have used dynamic culture conditions to increase EV production [12, 46, 48–50]. It is important to note that TEM images revealed no significant changes in morphology or size (**Figure 2E**). Furthermore, immunoblotting revealed no presence of the endoplasmic reticulum protein calnexin (**Figure 2F**), signifying that it is unlikely that the levels of shear stress used in this study were causing physical disruption to the cell membrane and/or instigating the release of apoptotic bodies.

MSCs monitor mechanical cues within their microenvironment in order to initiate adaptive and reparative responses; a capability governed by countless mechanosensors including, but not limited to, adherens junctions, focal adhesions, integrins, and ion channels [51–53]. Flow-derived shear stress is a mechanical cue pervasive throughout the physiological niche as well as many scalable manufacturing processes. However, flow and shear stress are two separate phenomena in that, besides inducing various stresses on the cell surface (i.e., shear, tensile, and compressive stresses), flow can also modulate the surrounding environment by augmenting diffusive exchanges (i.e., oxygen, nutrients, and waste metabolites) [54, 55]. After culturing the relatively mechanically-inert HEK293 cell line [33], we found a similar increase in both EV production and EV purity as was seen in MSCs (**Figure 3A, B**). HEK293 cells are often used in overexpression studies investigating mechanically-activated channels (e.g., Piezo2 and TRPV4) due to their low abundance of endogenous channels [33]. Coupled with our data, this could mean that the increased EV production we observed in this bioreactor system is a result of fluid dynamics rather than a specific mechanoresponse of the cells. However, in a recent study, while culturing MSCs in a flat-plate perfusion bioreactor, Kang et al. found that flow increased intracellular calcium as well as calcium-related proteins, and thus could be a mechanism behind the increase in EV biogenesis [14]. Separately, Guo and colleagues found that the increased MSC EV production present within their perfusion bioreactor was dependent on the expression of yes-associated protein (YAP), a well-known cellular mechanotransducer [12, 56, 57]. As both of these proposed mechanisms exist across cell-types (including HEK293), it would be of interest to probe these particular pathways within our system in future studies.

Of the various therapeutic applications of MSCs and MSC EVs elucidated thus far, the role they play in tissue reparative processes (e.g., angiogenesis) has been well-documented [58–61]. Moreover, it is known that flow-derived shear stress can be used to influence the regenerative properties of both MSCs and their secreted EVs [14]. Across two orthogonal angiogenic assays (i.e., cell migration and tube formation), we found that the angiogenic properties of MSC EVs derived from the perfusion bioreactor were at least as effective as those from flask-grown MSCs (**Figure 4B, C)**. We found that the same bioactivity pattern was apparent when using bone marrow-derived MSC EVs from several donors, but not when using MSC EVs derived from adipose tissue (**Figure S3**). This emphasizes the impact and importance of choosing the optimal EV source for the desired application [62]. The observed similarity in angiogenic activity between the flask- and bioreactor-derived MSC EVs were supported by similar cellular uptake of the EVs by HUVECs and comparable surface protein fingerprints (**Figure 6B, D**). Previous works have shown that cell culture conditions (e.g., bioreactor vs. flask or 2D vs. 3D culture) can enhance EV efficacy, alter EV uptake by recipient cells, and change the EV proteome profile [50, 63, 64]. Importantly, proteomic profiling of EVs utilizing the technique used in the current investigation (i.e., MALDI-TOF MS) can be sensitive. A study by Han and colleagues demonstrates the utility of this methodology by using the MALDI-TOF mass spectra of plasma-derived EVs to identify those patients with or without osteosarcoma lung metastasis [26]. While we reveal possible slight differences in the protein make-up (**Figure 6D**), we only investigated the surface proteins of undigested EV samples and thus further proteomic analyses (i.e., LC-MS) may reveal more significant differences. Additionally, it would be interesting to gauge how perfusion bioreactor culture can alter the proteome of MSC EVs in comparison with parental cells, which would be valuable information for the continued development of EVs in general for both diagnostic and therapeutic applications. It is also important to consider that any nuances caused by shear stress may alternatively lie within the EV metabolome, lipidome, and/or transcriptome, all of which are known to be dynamic with changing culture conditions [65, 66].

As described above, the flask- and bioreactor-derived MSC EVs were similar in numerous ways. However, there was a significant decrease in protein content per EV from the perfusion bioreactor. Consequently, dosing by protein entailed applying 5-7 times more bioreactor EVs (according to NTA data) than flask EVs. Taking this into account, we also dosed by EV count and found a similar pattern in the angiogenic activity assays, indicating a possible saturation point and equal efficacy of the two EV preparations (**Figure 4B**). This agrees with previous research showing that bioactivity of EVs can be maintained within bioreactor culture systems [18]. Critically, with the exaggerated EV production from the perfusion bioreactor, dosing by EV count allowed the application of a much smaller volume to achieve the same effect (e.g., 8 μl of bioreactor EVs vs 80 μl of flask EVs). Altogether, these results indicate that our perfusion bioreactor system may not enhance MSC EV efficacy but can be used to increase MSC EV potency and thus has important implications in scale-up efforts needed for clinical administration.

Intriguingly, when applied in a diabetic mouse wound healing model and compared to a vehicle control, EVs from the perfusion bioreactor system significantly improved healing in terms of percent wound closed and the number of new blood vessels formed, while those from flask culture had no effect (**Figure 5A, C**). The disparate results between the *in vitro* and *in vivo* data are not to be unexpected as the physiological niche is much more complex than anything that can be recapitulated in a laboratory setting. Particularly, wound healing is a highly regulated process that requires a dynamic yet balanced interplay among various cell types, including macrophages, fibroblasts, endothelial cells, and keratinocytes [58, 67]. Notably, MSC EVs are internalized by all of these cell types and have been documented to influence critical events during wound healing such as inflammation, collagen deposition, apoptosis, and re-epithelialization [58]. The *in vivo* results suggest that bioreactor MSC EVs may be more efficacious in any number of these wound healing processes and thus, further work is required to understand the exact role the MSC EVs play at the physiological wound site. It is also important to note that the bioreactor MSC EVs were able to alter the gene expression of HDMECs, although not significantly (**Figure 7B**). While the altered gene expression did not seem to impart any significant pro-angiogenic benefits in the *in vitro* validation assays, it remains to be seen how this translates to physiological wound healing.

Overall, the perfusion bioreactor utilized in the current study embodies a means to both enhance MSC EV potency while providing the opportunity to explore the effects of scale-up procedures necessary for clinical translation. The modularity of the system allows significant ability to tune the system to various needs, including the ability to alter other important attributes such as the stiffness and architecture of the 3D-printed scaffold. This investigation and further exploration of the system will ultimately inform the design of a manufacturing platform capable of producing a potent MSC EV wound healing therapeutic at the scale necessary for clinical use.

## Acknowledgements

This work was supported by the National Institutes of Health (HL141611, NS110637, GM130923, HL141922, HL159590 to SMJ; EB023833 to JPF) and the National Science Foundation (1750542 to SMJ; 1856350 to JPF). SMK, DL and LJB were supported by A. James Clark Doctoral Fellowships from the University of Maryland. DBP was supported by the American Heart Association (16PRE30770016). JWH was supported by the NIH (HL141611) and the Hendrix Burn/Wound Fund of Johns Hopkins University. The authors would like to thank Drs. Edward Eisenstein and James Parsons for technical advice and guidance in MALDI mass spectrometry.

## Data Availability Statement

The raw data required to reproduce these findings are available from the corresponding author upon reasonable request.

## Literature Cited

[1] M. Jafarinia, F. Alsahebfosoul, H. Salehi, N. Eskandari, M. Ganjalikhani-Hakemi, Mesenchymal Stem Cell-Derived Extracellular Vesicles: A Novel Cell-Free Therapy, Immunol Invest 49(7) (2020) 758–780.

[2] P. Vader, E.A. Mol, G. Pasterkamp, R.M. Schiffelers, Extracellular vesicles for drug delivery, Adv Drug Deliv Rev 106(Pt A) (2016) 148–156.

[3] O.M. Elsharkasy, J.Z. Nordin, D.W. Hagey, O.G. de Jong, R.M. Schiffelers, S.E. Andaloussi, P. Vader, Extracellular vesicles as drug delivery systems: Why and how?, Adv Drug Deliv Rev (2020).

[4] D. Levy, A. Jeyaram, L.J. Born, K.-H. Chang, S.N. Abadchi, A.T. Wei Hsu, T. Solomon, A. Aranda, S. Stewart, X. He, J.W. Harmon, S.M. Jay, The Impact of Storage Condition and Duration on Function of Native and Cargo-Loaded Mesenchymal Stromal Cell Extracellular Vesicles, BioRxiv (2022).

[5] X. Zhuang, X. Xiang, W. Grizzle, D. Sun, S. Zhang, R.C. Axtell, S. Ju, J. Mu, L. Zhang, L. Steinman, D. Miller, H.G. Zhang, Treatment of brain inflammatory diseases by delivering exosome encapsulated anti-inflammatory drugs from the nasal region to the brain, Mol Ther 19(10) (2011) 1769–79.

[6] R.O. Elliott, M. He, Unlocking the Power of Exosomes for Crossing Biological Barriers in Drug Delivery, Pharmaceutics 13(1) (2021).

[7] B. Crivelli, T. Chlapanidas, S. Perteghella, E. Lucarelli, L. Pascucci, A.T. Brini, I. Ferrero, M. Marazzi, A. Pessina, M.L. Torre, G. Italian Mesenchymal Stem Cell, Mesenchymal stem/stromal cell extracellular vesicles: From active principle to next generation drug delivery system, J Control Release 262 (2017) 104–117.

[8] L.J. Born, K.H. Chang, P. Shoureshi, F. Lay, S. Bengali, A.T.W. Hsu, S.N. Abadchi, J.W. Harmon, S.M. Jay, HOTAIR-Loaded Mesenchymal Stem/Stromal Cell Extracellular Vesicles Enhance Angiogenesis and Wound Healing, Adv Healthc Mater (2021) e2002070.

[9] E. Ragni, C. Perucca Orfei, P. De Luca, C. Mondadori, M. Vigano, A. Colombini, L. de Girolamo, Inflammatory priming enhances mesenchymal stromal cell secretome potential as a clinical product for regenerative medicine approaches through secreted factors and EV-miRNAs: the example of joint disease, Stem Cell Res Ther 11(1) (2020) 165.

[10] S. Ding, P. Kingshott, H. Thissen, M. Pera, P.Y. Wang, Modulation of human mesenchymal and pluripotent stem cell behavior using biophysical and biochemical cues: A review, Biotechnol Bioeng 114(2) (2017) 260–280.

[11] J. Gobin, G. Muradia, J. Mehic, C. Westwood, L. Couvrette, A. Stalker, S. Bigelow, C.C. Luebbert, F.S. Bissonnette, M.J.W. Johnston, S. Sauve, R.Y. Tam, L. Wang, M. Rosu-Myles, J.R. Lavoie, Hollow-fiber bioreactor production of extracellular vesicles from human bone marrow mesenchymal stromal cells yields nanovesicles that mirrors the immuno-modulatory antigenic signature of the producer cell, Stem Cell Res Ther 12(1) (2021) 127.

[12] S. Guo, L. Debbi, B. Zohar, R. Samuel, R.S. Arzi, A.I. Fried, T. Carmon, D. Shevach, I. Redenski, I. Schlachet, A. Sosnik, S. Levenberg, Stimulating Extracellular Vesicles Production from Engineered Tissues by Mechanical Forces, Nano Lett 21(6) (2021) 2497–2504.

[13] S. Morelli, A. Piscioneri, S. Salerno, L. De Bartolo, Hollow Fiber and Nanofiber Membranes in Bioartificial Liver and Neuronal Tissue Engineering, Cells Tissues Organs (2021) 1–30.

[14] H. Kang, Y.H. Bae, Y. Kwon, S. Kim, J. Park, Extracellular Vesicles Generated Using Bioreactors and their Therapeutic Effect on the Acute Kidney Injury Model, Adv Healthc Mater 11(4) (2022) e2101606.

[15] J. An, J.E.M. Teoh, R. Suntornnond, C.K. Chua, Design and 3D Printing of Scaffolds and Tissues, Engineering 1(2) (2015) 261–268.

[16] M. Harmati, E. Gyukity-Sebestyen, G. Dobra, L. Janovak, I. Dekany, O. Saydam, E. Hunyadi-Gulyas, I. Nagy, A. Farkas, T. Pankotai, Z. Ujfaludi, P. Horvath, F. Piccinini, M. Kovacs, T. Biro, K. Buzas, Small extracellular vesicles convey the stress-induced adaptive responses of melanoma cells, Sci Rep 9(1) (2019) 15329.

[17] O.G. de Jong, M.C. Verhaar, Y. Chen, P. Vader, H. Gremmels, G. Posthuma, R.M. Schiffelers, M. Gucek, B.W. van Balkom, Cellular stress conditions are reflected in the protein and RNA content of endothelial cell-derived exosomes, J Extracell Vesicles 1 (2012).

[18] D.B. Patel, C.R. Luthers, M.J. Lerman, J.P. Fisher, S.M. Jay, Enhanced extracellular vesicle production and ethanol-mediated vascularization bioactivity via a 3D-printed scaffold-perfusion bioreactor system, Acta Biomater 95 (2019) 236–244.

[19] T.N. Lamichhane, C.A. Leung, L.Y. Douti, S.M. Jay, Ethanol Induces Enhanced Vascularization Bioactivity of Endothelial Cell-Derived Extracellular Vesicles via Regulation of MicroRNAs and Long Non-Coding RNAs, Sci Rep 7(1) (2017) 13794.

[20] M.F. Pittenger, D.E. Discher, B.M. Peault, D.G. Phinney, J.M. Hare, A.I. Caplan, Mesenchymal stem cell perspective: cell biology to clinical progress, NPJ Regen Med 4 (2019) 22.

[21] A. Casado-Diaz, J.M. Quesada-Gomez, G. Dorado, Extracellular Vesicles Derived From Mesenchymal Stem Cells (MSC) in Regenerative Medicine: Applications in Skin Wound Healing, Front Bioeng Biotechnol 8 (2020) 146.

[22] X. He, Z. Dong, Y. Cao, H. Wang, S. Liu, L. Liao, Y. Jin, L. Yuan, B. Li, MSC-Derived Exosome Promotes M2 Polarization and Enhances Cutaneous Wound Healing, Stem Cells Int 2019 (2019) 7132708.

[23] T.N. Lamichhane, A. Jeyaram, D.B. Patel, B. Parajuli, N.K. Livingston, N. Arumugasaamy, J.S. Schardt, S.M. Jay, Oncogene Knockdown via Active Loading of Small RNAs into Extracellular Vesicles by Sonication, Cell Mol Bioeng 9(3) (2016) 315–324.

[24] T.N. Lamichhane, R.S. Raiker, S.M. Jay, Exogenous DNA Loading into Extracellular Vesicles via Electroporation is Size-Dependent and Enables Limited Gene Delivery, Mol Pharm 12(10) (2015) 3650–7.

[25] D.B. Patel, K.M. Gray, Y. Santharam, T.N. Lamichhane, K.M. Stroka, S.M. Jay, Impact of cell culture parameters on production and vascularization bioactivity of mesenchymal stem cell-derived extracellular vesicles, Bioeng Transl Med 2(2) (2017) 170–179.

[26] Z. Han, C. Peng, J. Yi, Y. Wang, Q. Liu, Y. Yang, S. Long, L. Qiao, Y. Shen, Matrix-assisted laser desorption ionization mass spectrometry profiling of plasma exosomes evaluates osteosarcoma metastasis, iScience 24(8) (2021) 102906.

[27] Y. Huang, J.Y. Qian, H. Cheng, X.M. Li, Effects of shear stress on differentiation of stem cells into endothelial cells, World J Stem Cells 13(7) (2021) 894–913.

[28] K.M. Kim, Y.J. Choi, J.H. Hwang, A.R. Kim, H.J. Cho, E.S. Hwang, J.Y. Park, S.H. Lee, J.H. Hong, Shear stress induced by an interstitial level of slow flow increases the osteogenic differentiation of mesenchymal stem cells through TAZ activation, PLoS One 9(3) (2014) e92427.

[29] A.C. Shieh, M.A. Swartz, Regulation of tumor invasion by interstitial fluid flow, Phys Biol 8(1) (2011) 015012.

[30] S. El Andaloussi, I. Mäger, X.O. Breakefield, M.J.A. Wood, Extracellular vesicles: biology and emerging therapeutic opportunities, Nature Reviews Drug Discovery 12 (2013) 347.

[31] A. Taherian, X. Li, Y. Liu, T.A. Haas, Differences in integrin expression and signaling within human breast cancer cells, BMC Cancer 11 (2011) 293.

[32] C. Alcaino, K. Knutson, P.A. Gottlieb, G. Farrugia, A. Beyder, Mechanosensitive ion channel Piezo2 is inhibited by D-GsMTx4, Channels (Austin) 11(3) (2017) 245–253.

[33] R. Soffe, S. Baratchi, S.Y. Tang, M. Nasabi, P. McIntyre, A. Mitchell, K. Khoshmanesh, Analysing calcium signalling of cells under high shear flows using discontinuous dielectrophoresis, Sci Rep 5 (2015) 11973.

[34] S. Palchetti, V. Colapicchioni, L. Digiacomo, G. Caracciolo, D. Pozzi, A.L. Capriotti, G. La Barbera, A. Lagana, The protein corona of circulating PEGylated liposomes, Biochim Biophys Acta 1858(2) (2016) 189–96.

[35] D.T. Jayaram, S.M. Pustulka, R.G. Mannino, W.A. Lam, C.K. Payne, Protein Corona in Response to Flow: Effect on Protein Concentration and Structure, Biophys J 115(2) (2018) 209–216.

[36] J. Phan, P. Kumar, D. Hao, K. Gao, D. Farmer, A. Wang, Engineering mesenchymal stem cells to improve their exosome efficacy and yield for cell-free therapy, J Extracell Vesicles 7(1) (2018) 1522236.

[37] M. Maumus, P. Rozier, J. Boulestreau, C. Jorgensen, D. Noel, Mesenchymal Stem Cell-Derived Extracellular Vesicles: Opportunities and Challenges for Clinical Translation, Front Bioeng Biotechnol 8 (2020) 997.

[38] M. Mendt, S. Kamerkar, H. Sugimoto, K.M. McAndrews, C.C. Wu, M. Gagea, S. Yang, E.V.R. Blanko, Q. Peng, X. Ma, J.R. Marszalek, A. Maitra, C. Yee, K. Rezvani, E. Shpall, V.S. LeBleu, R. Kalluri, Generation and testing of clinical-grade exosomes for pancreatic cancer, JCI Insight 3(8) (2018).

[39] D.E. Rodriguez-Fuentes, L.E. Fernandez-Garza, J.A. Samia-Meza, S.A. Barrera-Barrera, A.I. Caplan, H.A. Barrera-Saldana, Mesenchymal Stem Cells Current Clinical Applications: A Systematic Review, Arch Med Res 52(1) (2021) 93–101.

[40] J. Lembong, M.J. Lerman, T.J. Kingsbury, C.I. Civin, J.P. Fisher, A Fluidic Culture Platform for Spatially Patterned Cell Growth, Differentiation, and Cocultures, Tissue Eng Part A 24(23-24) (2018) 1715–1732.

[41] B.A. Gonzalez, M. Perez-Nevarez, A. Mirza, M.G. Perez, Y.M. Lin, C.D. Hsu, A. Caobi, A. Raymond, M.E. Gomez Hernandez, F. Fernandez-Lima, F. George, S. Ramaswamy, Physiologically Relevant Fluid-Induced Oscillatory Shear Stress Stimulation of Mesenchymal Stem Cells Enhances the Engineered Valve Matrix Phenotype, Front Cardiovasc Med 7 (2020) 69.

[42] C. Thery, K.W. Witwer, E. Aikawa, M.J. Alcaraz, J.D. Anderson, R. Andriantsitohaina, A. Antoniou, T. Arab, F. Archer, G.K. Atkin-Smith, D.C. Ayre, J.M. Bach, D. Bachurski, H. Baharvand, L. Balaj, S. Baldacchino, N.N. Bauer, A.A. Baxter, M. Bebawy, C. Beckham, A. Bedina Zavec, A. Benmoussa, A.C. Berardi, P. Bergese, E. Bielska, C. Blenkiron, S. Bobis-Wozowicz, E. Boilard, W. Boireau, A. Bongiovanni, F.E. Borras, S. Bosch, C.M. Boulanger, X. Breakefield, A.M. Breglio, M.A. Brennan, D.R. Brigstock, A. Brisson, M.L. Broekman, J.F. Bromberg, P. Bryl-Gorecka, S. Buch, A.H. Buck, D. Burger, S. Busatto, D. Buschmann, B. Bussolati, E.I. Buzas, J.B. Byrd, G. Camussi, D.R. Carter, S. Caruso, L.W. Chamley, Y.T. Chang, C. Chen, S. Chen, L. Cheng, A.R. Chin, A. Clayton, S.P. Clerici, A. Cocks, E. Cocucci, R. J. Coffey, A. Cordeiro-da-Silva, Y. Couch, F.A. Coumans, B. Coyle, R. Crescitelli, M.F. Criado, C. D’Souza-Schorey, S. Das, A. Datta Chaudhuri, P. de Candia, E.F. De Santana, O. De Wever, H.A. Del Portillo, T. Demaret, S. Deville, A. Devitt, B. Dhondt, D. Di Vizio, L.C. Dieterich, V. Dolo, A.P. Dominguez Rubio, M. Dominici, M.R. Dourado, T.A. Driedonks, F.V. Duarte, H.M. Duncan, R.M. Eichenberger, K. Ekstrom, S. El Andaloussi, C. Elie-Caille, U. Erdbrugger, J.M. Falcon-Perez, F. Fatima, J.E. Fish, M. Flores-Bellver, A. Forsonits, A. Frelet-Barrand, F. Fricke, G. Fuhrmann, S. Gabrielsson, A. Gamez-Valero, C. Gardiner, K. Gartner, R. Gaudin, Y.S. Gho, B. Giebel, C. Gilbert, M. Gimona, I. Giusti, D.C. Goberdhan, A. Gorgens, S. M. Gorski, D.W. Greening, J.C. Gross, A. Gualerzi, G.N. Gupta, D. Gustafson, A. Handberg, R. A. Haraszti, P. Harrison, H. Hegyesi, A. Hendrix, A.F. Hill, F.H. Hochberg, K.F. Hoffmann, B. Holder, H. Holthofer, B. Hosseinkhani, G. Hu, Y. Huang, V. Huber, S. Hunt, A.G. Ibrahim, T. Ikezu, J.M. Inal, M. Isin, A. Ivanova, H.K. Jackson, S. Jacobsen, S.M. Jay, M. Jayachandran, G. Jenster, L. Jiang, S.M. Johnson, J.C. Jones, A. Jong, T. Jovanovic-Talisman, S. Jung, R. Kalluri, S. I. Kano, S. Kaur, Y. Kawamura, E.T. Keller, D. Khamari, E. Khomyakova, A. Khvorova, P. Kierulf, K.P. Kim, T. Kislinger, M. Klingeborn, D.J. Klinke, 2nd, M. Kornek, M.M. Kosanovic, A.F. Kovacs, E.M. Kramer-Albers, S. Krasemann, M. Krause, I.V. Kurochkin, G.D. Kusuma, S. Kuypers, S. Laitinen, S.M. Langevin, L.R. Languino, J. Lannigan, C. Lasser, L.C. Laurent, G. Lavieu, E. Lazaro-Ibanez, S. Le Lay, M.S. Lee, Y.X.F. Lee, D.S. Lemos, M. Lenassi, A. Leszczynska, I.T. Li, K. Liao, S.F. Libregts, E. Ligeti, R. Lim, S.K. Lim, A. Line, K. Linnemannstons, A. Llorente, C.A. Lombard, M.J. Lorenowicz, A.M. Lorincz, J. Lotvall, J. Lovett, M.C. Lowry, X. Loyer, Q. Lu, B. Lukomska, T.R. Lunavat, S.L. Maas, H. Malhi, A. Marcilla, J. Mariani, J. Mariscal, E.S. Martens-Uzunova, L. Martin-Jaular, M.C. Martinez, V.R. Martins, M. Mathieu, S. Mathivanan, M. Maugeri, L.K. McGinnis, M.J. McVey, D.G. Meckes, Jr., K.L. Meehan, I. Mertens, V.R. Minciacchi, A. Moller, M. Moller Jorgensen, A. Morales-Kastresana, J. Morhayim, F. Mullier, M. Muraca, L. Musante, V. Mussack, D.C. Muth, K.H. Myburgh, T. Najrana, M. Nawaz, I. Nazarenko, P. Nejsum, C. Neri, T. Neri, R. Nieuwland, L. Nimrichter, J.P. Nolan, E.N. Nolte-’t Hoen, N. Noren Hooten, L. O’Driscoll, T. O’Grady, A. O’Loghlen, T. Ochiya, M. Olivier, A. Ortiz, L.A. Ortiz, X. Osteikoetxea, O. Ostergaard, M. Ostrowski, J. Park, D.M. Pegtel, H. Peinado, F. Perut, M.W. Pfaffl, D.G. Phinney, B.C. Pieters, R. C. Pink, D.S. Pisetsky, E. Pogge von Strandmann, I. Polakovicova, I.K. Poon, B.H. Powell, I. Prada, L. Pulliam, P. Quesenberry, A. Radeghieri, R.L. Raffai, S. Raimondo, J. Rak, M.I. Ramirez, G. Raposo, M.S. Rayyan, N. Regev-Rudzki, F.L. Ricklefs, P.D. Robbins, D.D. Roberts, S.C. Rodrigues, E. Rohde, S. Rome, K.M. Rouschop, A. Rughetti, A.E. Russell, P. Saa, S. Sahoo, E. Salas-Huenuleo, C. Sanchez, J.A. Saugstad, M.J. Saul, R.M. Schiffelers, R. Schneider, T.H. Schoyen, A. Scott, E. Shahaj, S. Sharma, O. Shatnyeva, F. Shekari, G.V. Shelke, A. K. Shetty, K. Shiba, P.R. Siljander, A.M. Silva, A. Skowronek, O.L. Snyder, 2nd, R.P. Soares, B. W. Sodar, C. Soekmadji, J. Sotillo, P.D. Stahl, W. Stoorvogel, S.L. Stott, E.F. Strasser, S. Swift, H. Tahara, M. Tewari, K. Timms, S. Tiwari, R. Tixeira, M. Tkach, W.S. Toh, R. Tomasini, A.C. Torrecilhas, J.P. Tosar, V. Toxavidis, L. Urbanelli, P. Vader, B.W. van Balkom, S.G. van der Grein, J. Van Deun, M.J. van Herwijnen, K. Van Keuren-Jensen, G. van Niel, M.E. van Royen, A.J. van Wijnen, M.H. Vasconcelos, I.J. Vechetti, Jr., T.D. Veit, L.J. Vella, E. Velot, F.J. Verweij, B. Vestad, J.L. Vinas, T. Visnovitz, K.V. Vukman, J. Wahlgren, D.C. Watson, M.H. Wauben, A. Weaver, J.P. Webber, V. Weber, A.M. Wehman, D.J. Weiss, J.A. Welsh, S. Wendt, A.M. Wheelock, Z. Wiener, L. Witte, J. Wolfram, A. Xagorari, P. Xander, J. Xu, X. Yan, M. Yanez-Mo, H. Yin, Y. Yuana, V. Zappulli, J. Zarubova, V. Zekas, J.Y. Zhang, Z. Zhao, L. Zheng, A.R. Zheutlin, A.M. Zickler, P. Zimmermann, A.M. Zivkovic, D. Zocco, E.K. Zuba-Surma, Minimal information for studies of extracellular vesicles 2018 (MISEV2018): a position statement of the International Society for Extracellular Vesicles and update of the MISEV2014 guidelines, J Extracell Vesicles 7(1) (2018) 1535750.

[43] J. Webber, A. Clayton, How pure are your vesicles?, J Extracell Vesicles 2 (2013).

[44] G. Raposo, W. Stoorvogel, Extracellular vesicles: exosomes, microvesicles, and friends, J Cell Biol 200(4) (2013) 373–83.

[45] Z. Andreu, M. Yanez-Mo, Tetraspanins in extracellular vesicle formation and function, Front Immunol 5 (2014) 442.

[46] M. Piffoux, A. Nicolas-Boluda, V. Mulens-Arias, S. Richard, G. Rahmi, F. Gazeau, C. Wilhelm, A.K.A. Silva, Extracellular vesicles for personalized medicine: The input of physically triggered production, loading and theranostic properties, Adv Drug Deliv Rev 138 (2019) 247–258.

[47] N.C. Gauthier, M.A. Fardin, P. Roca-Cusachs, M.P. Sheetz, Temporary increase in plasma membrane tension coordinates the activation of exocytosis and contraction during cell spreading, Proc Natl Acad Sci U S A 108(35) (2011) 14467–72.

[48] A.E. Morrell, G.N. Brown, S.T. Robinson, R.L. Sattler, A.D. Baik, G. Zhen, X. Cao, L.F. Bonewald, W. Jin, L.C. Kam, X.E. Guo, Mechanically induced Ca(2+) oscillations in osteocytes release extracellular vesicles and enhance bone formation, Bone Res 6 (2018) 6.

[49] D.C. Watson, D. Bayik, A. Srivatsan, C. Bergamaschi, A. Valentin, G. Niu, J. Bear, M. Monninger, M. Sun, A. Morales-Kastresana, J.C. Jones, B.K. Felber, X. Chen, I. Gursel, G.N. Pavlakis, Efficient production and enhanced tumor delivery of engineered extracellular vesicles, Biomaterials 105 (2016) 195–205.

[50] J. Cao, B. Wang, T. Tang, L. Lv, Z. Ding, Z. Li, R. Hu, Q. Wei, A. Shen, Y. Fu, B. Liu, Three-dimensional culture of MSCs produces exosomes with improved yield and enhanced therapeutic efficacy for cisplatin-induced acute kidney injury, Stem Cell Res Ther 11(1) (2020) 206.

[51] C.M. Potter, K.H. Lao, L. Zeng, Q. Xu, Role of biomechanical forces in stem cell vascular lineage differentiation, Arterioscler Thromb Vasc Biol 34(10) (2014) 2184–90.

[52] S.M. Naqvi, L.M. McNamara, Stem Cell Mechanobiology and the Role of Biomaterials in Governing Mechanotransduction and Matrix Production for Tissue Regeneration, Front Bioeng Biotechnol 8 (2020) 597661.

[53] N. Raman, S.A.M. Imran, K.B. Ahmad Amin Noordin, W. Zaman, F. Nordin, Mechanotransduction in Mesenchymal Stem Cells (MSCs) Differentiation: A Review, Int J Mol Sci 23(9) (2022).

[54] T.L. Place, F.E. Domann, A.J. Case, Limitations of oxygen delivery to cells in culture: An underappreciated problem in basic and translational research, Free Radic Biol Med 113 (2017) 311–322.

[55] J. Lovecchio, P. Gargiulo, J.L. Vargas Luna, E. Giordano, O.E. Sigurjonsson, A standalone bioreactor system to deliver compressive load under perfusion flow to hBMSC-seeded 3D chitosan-graphene templates, Sci Rep 9(1) (2019) 16854.

[56] S. Dupont, L. Morsut, M. Aragona, E. Enzo, S. Giulitti, M. Cordenonsi, F. Zanconato, J. Le Digabel, M. Forcato, S. Bicciato, N. Elvassore, S. Piccolo, Role of YAP/TAZ in mechanotransduction, Nature 474(7350) (2011) 179–83.

[57] X. Cai, K.C. Wang, Z. Meng, Mechanoregulation of YAP and TAZ in Cellular Homeostasis and Disease Progression, Front Cell Dev Biol 9 (2021) 673599.

[58] Y. Hou, J. Li, S. Guan, F. Witte, The therapeutic potential of MSC-EVs as a bioactive material for wound healing, Engineered Regeneration 2 (2021) 182–194.

[59] Q.L. Zeng, D.W. Liu, Mesenchymal stem cell-derived exosomes: An emerging therapeutic strategy for normal and chronic wound healing, World J Clin Cases 9(22) (2021) 6218–6233.

[60] C. Merino-Gonzalez, F.A. Zuniga, C. Escudero, V. Ormazabal, C. Reyes, E. Nova-Lamperti, C. Salomon, C. Aguayo, Mesenchymal Stem Cell-Derived Extracellular Vesicles Promote Angiogenesis: Potencial Clinical Application, Front Physiol 7 (2016) 24.

[61] E.R. Bray, R.S. Kirsner, E.V. Badiavas, Mesenchymal Stem Cell-Derived Extracellular Vesicles as an Advanced Therapy for Chronic Wounds, Cold Spring Harb Perspect Biol (2022).

[62] C. Almeria, S. Kress, V. Weber, D. Egger, C. Kasper, Heterogeneity of mesenchymal stem cell-derived extracellular vesicles is highly impacted by the tissue/cell source and culture conditions, Cell Biosci 12(1) (2022) 51.

[63] G.D. Kusuma, A. Li, D. Zhu, H. McDonald, I.M. Inocencio, D.C. Chambers, K. Sinclair, H. Fang, D.W. Greening, J.E. Frith, R. Lim, Effect of 2D and 3D Culture Microenvironments on Mesenchymal Stem Cell-Derived Extracellular Vesicles Potencies, Front Cell Dev Biol 10 (2022) 819726.

[64] S. Rocha, J. Carvalho, P. Oliveira, M. Voglstaetter, D. Schvartz, A.R. Thomsen, N. Walter, R. Khanduri, J.C. Sanchez, A. Keller, C. Oliveira, I. Nazarenko, 3D Cellular Architecture Affects MicroRNA and Protein Cargo of Extracellular Vesicles, Adv Sci (Weinh) 6(4) (2019) 1800948.

[65] M. Palviainen, H. Saari, O. Karkkainen, J. Pekkinen, S. Auriola, M. Yliperttula, M. Puhka, K. Hanhineva, P.R. Siljander, Metabolic signature of extracellular vesicles depends on the cell culture conditions, J Extracell Vesicles 8(1) (2019) 1596669.

[66] S. Thippabhotla, C. Zhong, M. He, 3D cell culture stimulates the secretion of in vivo like extracellular vesicles, Sci Rep 9(1) (2019) 13012.

[67] M. Rodrigues, N. Kosaric, C.A. Bonham, G.C. Gurtner, Wound Healing: A Cellular Perspective, Physiol Rev 99(1) (2019) 665–706.

